# Polyphyletic domestication and inter-lineage hybridization magnified genetic diversity of cultivated melon, *Cucumis melo* L

**DOI:** 10.1101/2024.06.27.601017

**Authors:** Katsunori Tanaka, Gentaro Shigita, Tran Phuong Dung, Phan Thi Phuong Nhi, Mami Takahashi, Yuki Monden, Hidetaka Nishida, Ryuji Ishikawa, Kenji Kato

**Author notes:** Corresponding author: Kenji Kato, Graduate School of Environmental, Life, Natural Science and Technology, Okayama University, 1-1-1 Tsushima Naka, Kita-ku, Okayama, 700-8530 Japan.

## Abstract

A total of 212 melon accessions with diverse geographical origins were classified into large and small seed-types by length of seed at the boundary of 9 mm, and into five populations based on polymorphisms in the nuclear genome. They were further divided into three maternal lineages, Ia, Ib, and Ic, by polymorphisms in the chloroplast genome. By combining these three classifications, the Europe/US subsp. *melo* and the East Asian subsp. *agrestis* were characterized as [large seed, Ib, PopA1 or A2] and [small seed, Ia, PopB1 or B2], respectively, indicating nearly perfect divergence in both nuclear and cytoplasm genomes. In contrast, in South and Southeast Asia, in addition to the Europe/US and East Asian types, recombinant types were also frequently found, indicating unclear genetic differentiation in South and Southeast Asia. Such an intermixed structure of genetic variation supported the Indian origin of Ia and Ib types of melon. Seed length was intermediate, between the large and small seed-types, and chloroplast type was a mixture of Ia and Ib in Momordica, suggesting its origin from the recombinant type. In Africa, three lineages of melon were distributed allopatrically and showed distinct divergence. Subsp. *agrestis* of the Ic type proved to be endemic to Africa, indicating its African origin.

## Introduction

Melon (*Cucumis melo* L.) is cultivated worldwide. Its history of cultivation can be dated back to the 3rd–2nd millennia BCE in Africa and the Asian continent (Zohary and Hopf 1988; Walters 1989; Kirkbride 1993). Reflecting this long history of cultivation, diverse types of cultivated melon have been established in various parts of the world, and melon is considered the most diversified species of Cucurbitaceae. It is separated into two subspecies, *melo* L. and *agrestis* Naudin (Naudin 1859), the former being typically grown in Europe and the US and the latter in East and South Asia. As summarized by Pitrat (2016), *C. melo* is divided into 19 horticultural groups based primarily on flower and fruit traits: Agrestis, Kachri, Chito, Tibish, Acidulus, Momordica, Conomon, Makuwa, Chinensis, Flexuosus, Chate, Dudaim, Chandalak, Indicus, Ameri, Cassaba, Ibericus, Inodorus, and Cantalupensis.

Previous studies of genetic diversity among melon accessions from diverse geographical origins highlighted three points concerning the evolution and diversification of melon, though the direct ancestral wild species has yet to be identified. The first point is the genetic divergence between Europe/US melons, Cantalupensis and Inodorus, and East Asian melons, Conomon and Makuwa, as demonstrated by analysis of seed size (Fujishita 1983; Sabato et al. 2015; Tanaka et al. 2016), fruit characteristics (Liu et al. 2004), molecular markers (Esteras et al. 2013), genome-wide Single Nucleotide Polymorphisms (SNPs) (Zhao et al. 2019; Liu et al. 2020; Wang et al. 2021; Shigita et al. 2023) and both phenotypic traits and molecular markers (Stepansky et al. 1999; Tanaka et al. 2013). The Europe/US melon is characterized by sweetness of the fruit flesh and seeds longer than 9.0 mm, the East Asian melon by less sweetness of the fruit flesh and seeds shorter than 9.0 mm. Molecular analysis of the nuclear genome also revealed distinct genetic differentiation between these two geographical groups of melon. Sequence polymorphisms of the chloroplast genome, including SNPs, In/Del, and Simple Sequence Repeats (SSRs), were detected by Tanaka et al. (2013) and suggested that the Europe/US melon subsp. *melo* and the East Asian melon subsp. *agrestis* were established in different maternal lineages, Ib and Ia, respectively. This finding strongly supported the independent origin of cultivated melon (Pitrat 2013; Endl et al. 2018; Zhao et al. 2019). However, it has not been confirmed, using a diverse collection of melon landraces, whether the two geographical groups of melon are differentiated in a set of morphological traits (e.g., seed size) and nuclear and chloroplast genomes.

The second point is genetic diversification in Indian melon. India had been considered the secondary center of diversity of cultivated melon (Robinson and Decker-Walters 1997), and higher phenotypic and genetic variations have been reported (Akashi et al. 2002; McCreight et al. 2004; Dhillon et al. 2007, 2012; Tanaka et al. 2007; Haldhar et al. 2018). Analysis of the chloroplast genome and seed length provided new insights into the diversity in India. The Indian melon consists of two maternal lineages and is an admixture of the Europe/US type (large seed type with Ib cytoplasm), the East Asian type (small seed type with Ia cytoplasm), and recombinant types between those two types (large seed type with Ia cytoplasm and small seed type with Ib cytoplasm) (Tanaka et al. 2013). Recombinants could be established by hybridization between the two types, and might largely contribute to the enrichment of genetic diversity. Another possible explanation for the large genetic variation is that melon originated around India, as suggested by Sebastian et al. (2010), John et al. (2013), Endl et al. (2018), and Zhao et al. (2019). Indian melon landraces, including Momordica and Flexuosus, should be analyzed in detail, to elucidate the evolution and diversification of melon and the genetic mechanism for the enriched genetic diversity.

The third point concerns the African origin of melon. Africa is thought to be another center of melon domestication (Zohary and Hopf 1988; Kirkbride 1993; Pitrat 2008). Analyses using Random Amplified Polymorphic DNA (RAPD) and/or SSR markers revealed large genetic variation in African melon landraces (Mliki et al. 2001; López-Sesé et al. 2002; Serres-Giardi and Dogimont 2012). Landraces from Northern Africa are closely related to those in Europe/US (Mliki et al. 2001; Nakata et al. 2005), while those from South and Western Africa are closely related to those in India and East Asia (Monforte et al. 2003; Esteras et al. 2013). Diversity in African melon was also demonstrated by analysis of the chloroplast genome, and all three cytoplasm types, Ia, Ib and Ic, were detected in 29 landraces examined (Tanaka et al. 2013). The Ia and Ib types accord with those in East Asia and in Europe/US, respectively. In contrast, the Ic type was found only in Western and Southern Africa and was rather distantly related to the other two types, suggesting the independent origin of the Ic type in the third lineage in Africa. To validate spatial genetic differentiation in Africa and its correspondence with that on other continents, the genetic structure of a worldwide collection of melon should be examined in detail.

Here, we demonstrate the independent origin of cultivated melon in three maternal lineages, Ia, Ib and Ic, and that the two subspecies *agrestis* and *melo* were established and diversified in the former two lineages, respectively, based on analyses of seed size, nuclear genotype and cytoplasm type, using 212 melon accessions from various parts of the world. We also discuss the genetic mechanism for diversification in the Indian melon and the independent origin of cultivated melon in the third lineage in Africa.

## Materials and Methods

### Plant materials

A total of 212 accessions of melon (*Cucumis melo* L.) were analyzed in this study, selected from a wide geographical area of Africa, Asia, Europe and North America. They consisted of 31 accessions of Agrestis, 84 of six horticultural groups classified after Pitrat (2016), and 97 unclassified landraces. The geographical origin and horticultural group of each accession are given in Supplemental Table 1. Since distinct seed size variation within accession was observed in two accessions from India, PI 210541 and PI 124112, they were separated into two accessions. Two accessions of Japanese Cantalupensis and one accession of Chinese Honeydew were included in the Europe/US group for analysis, since they derived from recent introductions from Europe or the USA. Five accessions of the Chinese Hami melon were included in the West and Central Asian group, since it is distantly related to the Chinese Conomon and Makuwa and closely related to West and Central Asian melon (Aierken et al. 2011).

Seeds of these accessions were provided by the North Central Regional Plant Introduction Station, Iowa State University (USDA-ARS), USA; the NARO Institute of Vegetable and Floriculture Science (NIVFS), Japan; and Okayama University, Japan. These accessions were cultivated in the field or greenhouse of Okayama University, and self-pollinated to harvest mature seeds for size measurement.

### Seed size measurement

The lengths and widths of three representative seeds were measured for each accession. Based on seed length, each accession was classified into the large seed-type (≥ 9.0 mm) or small seed-type (< 9.0 mm) according to Akashi et al. (2002).

To confirm the adequacy of classification based on seed length using a broad spectrum of melon accessions, we obtained 100 seeds weight data of a total of 7,495 melon accessions from the USDA-ARS Germplasm Resources Information Network (GRIN). We then analyzed the relationship between seed length and 100 seeds weight by measuring the 100 seeds weights of 130 accessions included in this study (Supplemental Table 1). Based on the result, the 7,495 melon accessions were roughly classified into large and small seed-types at the boundary of 2.2 g of 100 seeds weight.

### DNA extraction

For each accession, genomic DNA was extracted from a single ten-day-old seedling, after Murray and Thompson (1980), with minor modifications.

### Chloroplast genotyping

Chloroplast genome type was determined by analysis of nine SNP markers, one In/Del marker and two SSR markers (Table 1). Of these 12 markers, ccSSR7 was developed by Chung and Staub (2003) and the others were newly developed in this study from sequence polymorphisms previously detected by Tanaka et al. (2013).

**Table 1.**
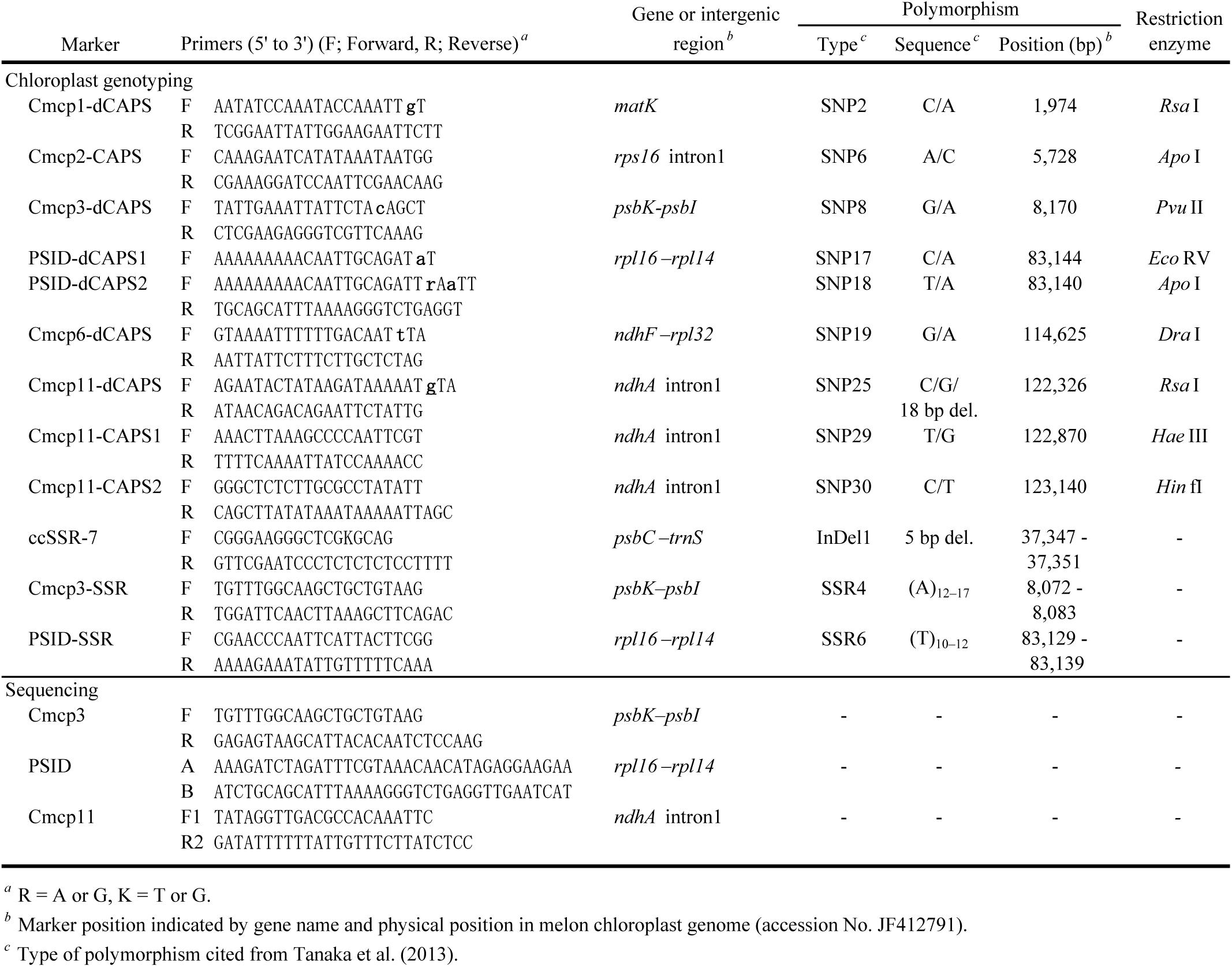
PCR primers used for genotyping the chloroplast genome and for sequencing target regions.

Nine SNPs were examined by three Cleaved Amplified Polymorphic Sequence (CAPS) and six derived CAPS (dCAPS) markers (Table 1). PCR amplification was done by using *Taq* DNA polymerase (Sigma-Aldrich^®^, USA), and the PCR product was digested with the restriction enzymes shown in Table 1. The digested product was electrophoresed in agarose gel for SNP genotyping. For PCR amplification of PSID-dCAPS1 and PSID-dCAPS2, PSID-R primer was used as a common reverse primer. For genotyping ccSSR-7, the deletion of 5 bp was detected by acrylamide gel electrophoresis according to Aierken et al. (2011).

For Cmcp3-SSR and PSID-SSR, PCR amplification was done using Ex*Taq*^TM^ DNA polymerase (Takara, Japan). The PCR product, including a fluorescently labeled size marker, was applied to a CEQ8000 capillary sequencer (Beckman Coulter, USA). The resulting chromatograms were visualized and analyzed using CEQ 8000 Fragment Analysis software.

Additional sequence polymorphisms were indicated by the analysis of Cmcp11-dCAPS, Cmcp3-SSR and PSID-SSR. The Cmcp11 region was sequenced for five accessions (PI 185111, 940101, 940103, 770134, and 770135), and Cmcp3 and PSID regions for three accessions (940099, 940102, and PI 614519). The experimental procedure is fully described by Tanaka et al. (2013), and the representative nucleotide sequences determined were registered in the DNA Data Bank of Japan (DDBJ) (Supplemental Table 2).

### RAPD analysis

RAPD analysis was carried out for 147 melon accessions. Eighteen random primers (12-mer, Bex, Japan) selected for their ability to detect polymorphism by Tanaka et al. (2007) were used to produce 27 markers (Supplemental Table 3). PCR amplification and electrophoresis were carried out according to Tanaka et al. (2007). For the remaining 65 accessions, RAPD data obtained by Tanaka et al. (2007) was used for data analysis.

### Data analysis

Tanaka et al. (2013) identified a total of 12 subtypes of chloroplast genome, and three additional subtypes, Ia-4, Ia-5, and Ia-6, were detected in this study. To investigate the relationship among 15 cytoplasm types, a Median-Joining (MJ) network (Bandelt et al. 1999) was constructed with the NETWORK program, v4.6.1.1. For the calculation, we used sequence data of 30 SNPs (SNP 1–30) and five In/Del (InDel 1–5) of three accessions sequenced in this study (940099, 940102, and PI 614519) and of 53 accessions including four wild species, *C*. *anguria*, *C*. *hystrix*, *C*. *metuliferus* and *C*. *sagittatus*, previously sequenced by Tanaka et al. (2013). The superfluous median vector on the constructed network tree was purged by the maximum parsimony option (Polzin and Daneschmand 2003). The same data set was used to construct neighbor-joining (NJ) and maximum-likelihood (ML) trees with a bootstrapping of 1,000 replications using MEGA v11 (Kumar et al. 2016)

Marker bands of RAPD were scored as 1 for a positive band and zero for a null band. The number of different alleles (Na), number of effective alleles (Ne), and expected heterozygosity (He) were calculated using GenALEX v6.503 (Peakall and Smouse 2006, 2012). The polymorphic information content (PIC) and gene diversity (*D*) within each group were then calculated according to Botstein et al. (1980) and Nei (1972). The genetic distance (GD) among accessions was calculated as described by Apostol et al. (1993). Based on the GD matrix, a dendrogram was constructed by using the Unweighted Pair Group Method with arithmetic mean (UPGMA), using PHYLIP v3.698 programs (https://evolution.genetics.washington.edu/phylip.html), and was compared with that by the NJ method. The relationships among melon groups were visualized by principal coordinate analysis (PCO) and quantified by fixation index (*F_ST_*) value by GenALEX v6.503. The model-based clustering program, STRUCTURE v2.3.4 (Pritchard et al. 2000), was used to infer population structure by a Bayesian approach from the RAPD marker data set. The optimal value of *K* (the number of clusters) was deduced by evaluating *K* = 1–10, and determined by an admixture model with an allele frequencies correlated model. The length of burn-in of the Markov Chain Monte Carlo (MCMC) iterations was set to 5,000 and data were collected over 5,000 MCMC iterations in each run. Twenty iterations per *K* were conducted. The optimal value of *K* was identified using the ad hoc procedure introduced by Pritchard et al. (2000) and the method developed by Evanno et al. (2005), which were carried out in Structure Harvester (Earl and vonHoldt 2012). Data plotting after the STRUCTURE simulation was conducted with CLUMPP (Jakobsson and Rosenberg 2007). Substructures within each major population were detected by repeating the simulation for each population with the same settings.

## Result

### Seed size measurement

Average seed length showed wide intraspecific variation, ranging from 3.92 mm to 15.09 mm among the 212 accessions, as shown in Fig. 1, and seed widths ranged from 1.80 mm to 7.00 mm. These two traits were closely correlated (*r* = 0.912, *p* < 0.01), and thus seed length was used as an indicator of seed size hereafter. The difference in seed length between geographical regions was significant (*p* < 0.01), though it was insignificant between Europe/US and West and Central Asia, and between South Asia and Southeast Asia. The analysis of seed length classified the Europe/US accessions into the large seed-type (≥ 9.0 mm) except for one Japanese netted accession, Melon Chuukanbohon Nou 1, bred by crossing Cantalupensis and Makuwa, while the East Asian accessions were classified as of small seed-type (< 9.0 mm). Among melon accessions from other regions, those from Northern Africa and West Asia were recognized as large seed-type. In contrast, the small seed-type predominated in Western and Southern Africa. These results demonstrate geographical differentiation in seed length.

**Fig. 1.**
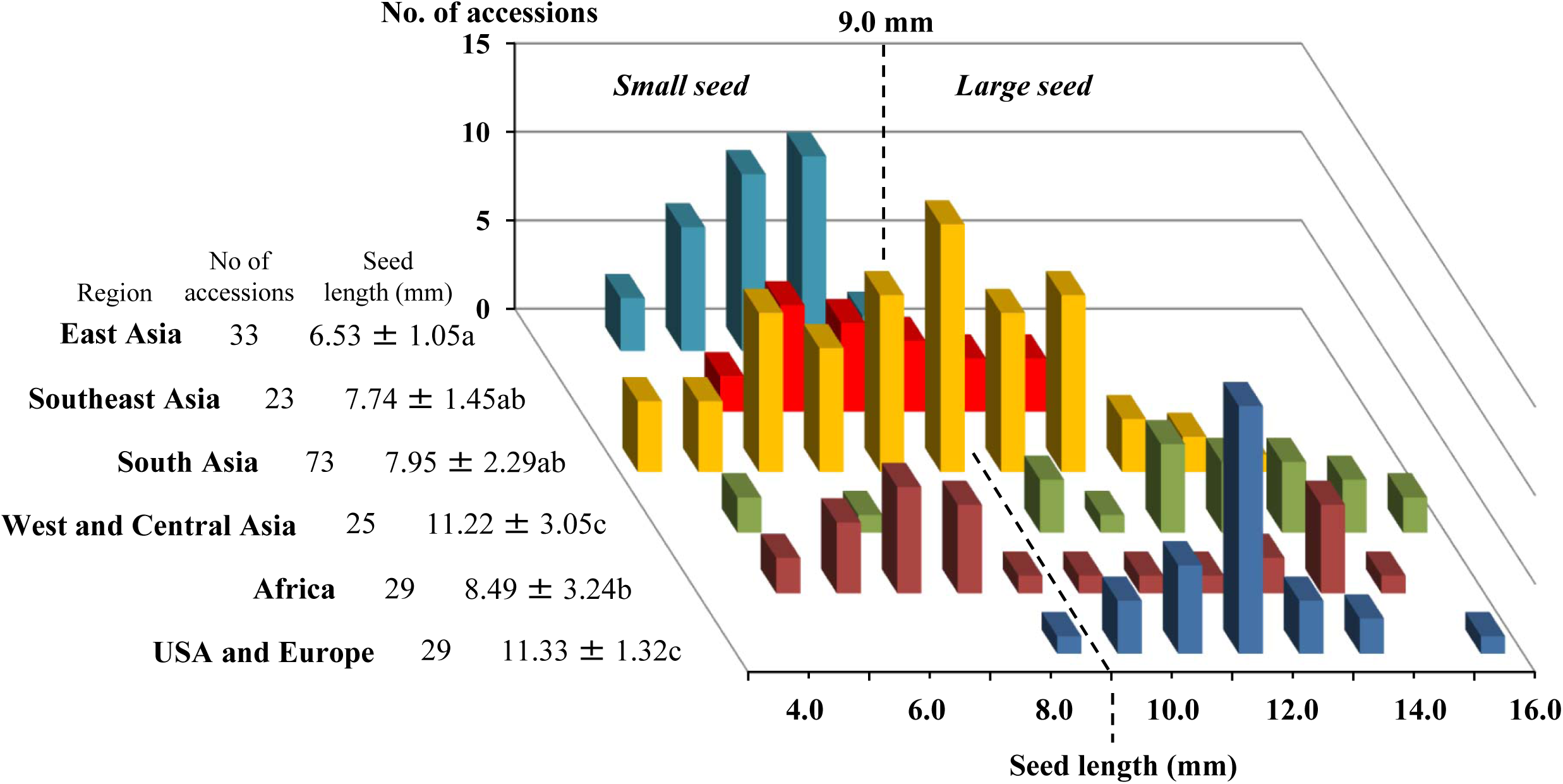
Frequency distribution of seed length in melon accessions from six geographical regions. Geographical differences were analysed by the Tukey-Kramer multiple comparison test with significant differences at p < 0.01.

Seed lengths showed continuous variation in South and Southeast Asia, where both large and small seed types were frequently found. In these regions, two classes (8–9 mm and 9–10 mm), flanking the boundary between the large and small seed-types, accounted for 30.2% of accessions, including 11 of the 22 accessions of Momordica and Flexuosus, whose average seed lengths were 9.3 mm and 9.7 mm, respectively. In contrast, the frequency of the two classes was low, being 6.1% in East Asia and 13.8% in Europe/US. Such a geographical difference was validated by the analysis of 7,495 melon accessions, whose seed weight was available instead of seed length. In order to classify these accessions into large and small seed types, we first investigated the relationship between seed length and 100 seeds weight in 130 accessions (Supplemental Table 1). The 100 seeds weights ranged from 2.03 g to 8.03 g (3.47 g on average) in 46 accessions of large seed-types, and from 0.36 g to 2.73 g (1.36 g on average) in 84 accessions of small seed-types (Supplemental Table 4). The difference in 100 seeds weights between the two seed types was significant (*p* < 0.01), and a highly positive correlation was observed between weight and length of seeds (*r* = 0.916, *p* < 0.01, Supplemental Fig. 3). From these results, the threshold to discriminate between the large and small seed-types was set at 2.2 g, by which 122 of 130 accessions (93.8%) were classified as respective seed length types. Based on these criteria, 1,998 accessions from India were separated into large and small seed-types, which accounted for 57.0% and 43.0%, respectively (Table 2). In addition, the proportions of two classes (1.8–2.2 g and 2.2–2.6 g) flanking the boundary between the large and small seed-types were high, being 19.6% and 14.6%, respectively, in good accordance with the result based on seed lengths.

**Table 2.** Geographical variation of 100 seeds weight among 7,495 accessions of melon.

| Area | Country | Total | 100 seeds weight (g) |  |  |  |  |  |  |  |  |  |  |  |  |  |  |  |  |  |  |  |  |  |  |  |  |
| --- | --- | --- | --- | --- | --- | --- | --- | --- | --- | --- | --- | --- | --- | --- | --- | --- | --- | --- | --- | --- | --- | --- | --- | --- | --- | --- | --- |
|  |  |  | -0.6 | -1.0 | -1.4 | -1.8 | -2.2 | -2.6 | -3.0 | -3.4 | -3.8 | -4.2 | -4.6 | -5.0 | -5.4 | -5.8 | -6.2 | -6.6 | -7.0 | -7.4 | -7.8 | -8.2 | -8.6 | -9.0 | -9.4 | -9.8 | 9.8 ≥ |
|  |  |  | Small seed |  |  |  |  | Large seed |  |  |  |  |  |  |  |  |  |  |  |  |  |  |  |  |  |  |  |
| USA | North | 224 | 40 | 8 | 4 | 6 | 20 | 26 | 52 | 33 | 21 | 6 | 5 | 1 | - | - | - | - | 1 | - | - | - | - | 1 | - | - | - |
|  | Caribbean | 27 | - | - | - | - | 1 | - | 2 | 4 | 7 | 5 | 6 | 1 | 1 | - | - | - | - | - | - | - | - | - | - | - | - |
|  | Central | 42 | 4 | 5 | 5 | 3 | - | 5 | 5 | 1 | 1 | 6 | 4 | 1 | 1 | - | - | - | - | 1 | - | - | - | - | - | - | - |
|  | South | 40 | 2 | 1 | - | - | - | 2 | 5 | 6 | 2 | 9 | 7 | 6 | - | - | - | - | - | - | - | - | - | - | - | - | - |
| Europe |  | 1577 | - | 1 | 4 | 16 | 22 | 53 | 89 | 179 | 205 | 248 | 221 | 207 | 125 | 90 | 55 | 28 | 20 | 6 | 4 | 2 | 1 | 1 | - | - | - |
| Russia /Former Soviet Union |  | 78 | - | 1 | - | 1 | 4 | 7 | 11 | 11 | 12 | 14 | 11 | - | 3 | - | 2 | 1 | - | - | - | - | - | - | - | - | - |
| Africa | North | 290 | - | 1 | - | 1 | 3 | 4 | 7 | 20 | 32 | 52 | 58 | 40 | 33 | 27 | 9 | 1 | 1 | - | - | - | - | - | - | - | 1 |
|  | East | 7 | - | - | - | - | - | 2 | - | - | 3 | 2 | - | - | - | - | - | - | - | - | - | - | - | - | - | - | - |
|  | West | 18 | 2 | 4 | 4 | - | - | - | 1 | 3 | 1 | 1 | 1 | 1 | - | - | - | - | - | - | - | - | - | - | - | - | - |
|  | South | 121 | 1 | 8 | 32 | 28 | 24 | 19 | 5 | 1 | 2 | 1 | - | - | - | - | - | - | - | - | - | - | - | - | - | - | - |
| West and Central Asia |  | 2615 | 9 | 3 | 11 | 25 | 44 | 100 | 157 | 254 | 351 | 389 | 413 | 310 | 246 | 153 | 78 | 33 | 18 | 13 | 2 | 4 | - | 1 | 1 | - | - |
| South Asia | Pakistan | 73 | - | 2 | - | 1 | 8 | 13 | 19 | 10 | 8 | 2 | 4 | 1 | 2 | 2 | - | - | - | 1 | - | - | - | - | - | - | - |
|  | India | 1998 | 57 | 158 | 199 | 332 | 392 | 291 | 220 | 146 | 82 | 79 | 29 | 9 | 4 | - | - | - | - | - | - | - | - | - | - | - | - |
|  | Maldives | 33 | 19 | 2 | - | 1 | 2 | 4 | 4 | 1 | - | - | - | - | - | - | - | - | - | - | - | - | - | - | - | - | - |
| Southeast Asia |  | 27 | 2 | 6 | 2 | 1 | 2 | 3 | 6 | 3 | 1 | 1 | - | - | - | - | - | - | - | - | - | - | - | - | - | - | - |
| East Asia |  | 325 | 2 | 36 | 93 | 55 | 15 | 13 | 21 | 15 | 22 | 20 | 10 | 7 | 9 | 4 | 1 | 2 | - | - | - | - | - | - | - | - | - |
| Total |  | 7495 | 138 | 236 | 354 | 470 | 537 | 542 | 604 | 687 | 750 | 835 | 769 | 584 | 424 | 276 | 145 | 65 | 40 | 21 | 6 | 6 | 1 | 3 | 1 | 0 | 1 |

### Chloroplast genome type

The sequence polymorphisms identified in the previous study (Tanaka et al. 2013) were detected by PCR analysis of CAPS, dCAPS, and SSR markers of the chloroplast genome (Fig. 2). As summarized in Table 3, new chloroplast genome subtypes Ia-4 to Ia-6 were detected in Agrestis from South Asia, with unique combinations of Cmcp11-dCAPS, Cmcp3-SSR and PSID-SSR alleles (Supplemental Table 1). In contrast, Ic-3 and Ic-4 could not be distinguished, since the diagnostic marker for SNP21 was not used in this study. Likewise, Ic-5 and Ic-6 were not distinguished, since the diagnostic marker for InDel2 was not used. Of the 212 accessions, 118, 82, and 12 were classified as Ia, Ib, and Ic types, respectively, and were further divided into 13 subtypes (Table 3). Five of the 118 accessions with Ia cytoplasm were classified as Ia-4 subtype, and one accession each as Ia-5 and Ia-6 subtypes.

**Fig. 2.**
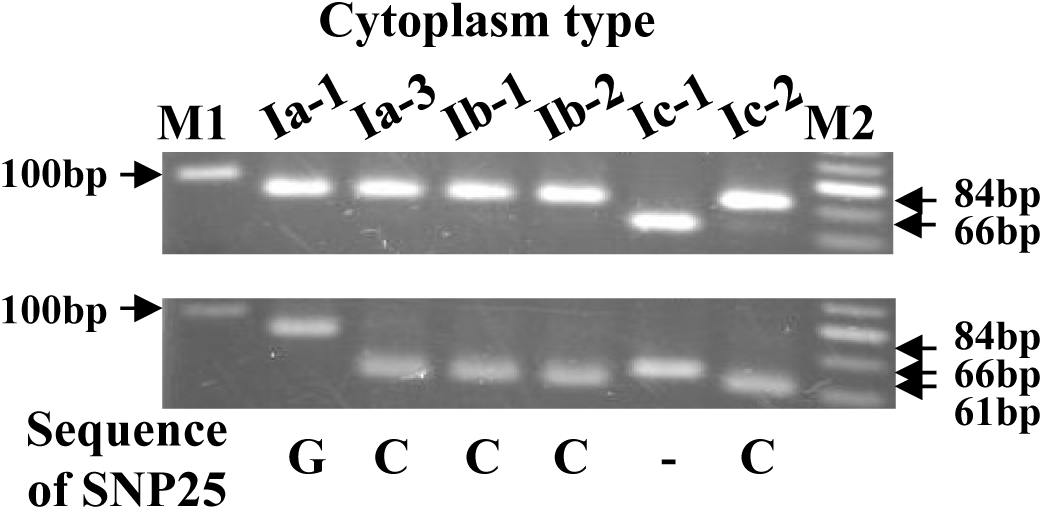
Gel image of dCAPS analysis of Cmcp11-dCAPS. Agarose gel electrophoresis patterns of amplified products (upper panel) and restriction fragments digested with *Rsa* I (lower panel). M1; 100 bp DNA Ladder (Takara, Japan), Ia-1; ‘Kinpyo’, Ia-3; PI 482424, Ib-1; 525105, Ib-2; ‘Earl’s Favourite’, Ic-1; PI 185111, Ic-2; PI 436533, M2; 20 bp DNA Ladder (Takara, Japan).

**Table 3.** Chloroplast genome types discriminated by the analysis of 12 diagnostic markers.

| Cytoplasm<br>type/<br>Species <sup>a</sup> | Number<br>of<br>accessions | SNP2<br>Cmcp1<br>-dCAPS | SNP6<br>Cmcp2<br>-CAPS | SNP8<br>Cmcp3<br>-dCAPS | SNP17<br>PSID-dCAPS1 | SNP18<br>PSID-dCAPS2 | SNP19<br>Cmcp6<br>-dCAPS | SNP25<br>Cmcp11<br>-dCAPS | SNP29<br>Cmcp11<br>-CAPS1 | SNP30<br>Cmcp11<br>-CAPS2 | InDel1<br>cc<br>SSR7 | SSR4<br>Cmcp3<br>-SSR2 | SSR6<br>PSID<br>-SSR |
| --- | --- | --- | --- | --- | --- | --- | --- | --- | --- | --- | --- | --- | --- |
| Ia-1 | 86 | C | C | G | A | A | A | G | G | T | - | (A)17 | (T)11 |
| Ia-2 | 2 | C | C | G | A | A | A | C | T | T | - | (A)17 | (T)11 |
| Ia-3 | 23 | C | C | G | A | A | A | C | T | T | - | (A)16 | (T)11 |
| Ia-4 | 5 | C | C | G | A | A | A | Deletion | T | T | - | (A)16 | (T)11 |
| Ia-5 | 1 | C | C | G | A | A | A | Deletion | T | T | - | (A)16 | (T)12 |
| Ia-6 | 1 | C | C | G | A | A | A | C | T | T | - | (A)15 | (T)11 |
| Ib-1 | 24 | A | C | A | A | T | G | C | T | T | - | (A)15 | (T)11 |
| Ib-2 | 36 | A | C | A | A | T | G | C | T | T | - | (A)15 | (T)10 |
| Ib-3 | 22 | A | C | A | A | T | G | C | T | T | Deletion | (A)15 | (T)11 |
| Ic-1 | 1 | C | A | G | C | T | G | Deletion | T | C | - | (A)13 | (T)11 |
| Ic-2 | 1 | C | A | G | C | T | G | C | T | C | - | (A)15 | (T)11 |
| Ic-3/-4 <sup>b</sup> | 4 | C | A | G | C | T | G | C | T | C | - | (A)14 | (T)11 |
| Ic-5/-6 <sup>c</sup> | 6 | C | A | G | C | T | G | C | T | C | - | (A)12 | (T)12 |
| *C. anguria* |  | C | A | G | A | T | G | Deletion | T | C | - | (A)14 | (T)11 |
| *C. hystrix* |  | C | A | G | C | T | G | Deletion | T | C | - | (A)13 | (T)11 |
| *C. metuliferus* |  | C | A | G | C | T | G | Deletion | T | C | - | (A)14 | (T)5 |
| *C. sagittatus* |  | C | A | G | C | T | Deletion | Deletion | T | C | - | (A)11 | (T)8 |
| Total | 212 |  |  |  |  |  |  |  |  |  |  |  |  |

<sup>a</sup> Sequence data was cited from Tanaka et al. (2013), with the exception of Ia-4, Ia-5 and Ia-6 which were sequenced in this study.
<sup>b</sup> Ic-3 and Ic-4 shows identical genotype, since SNP21 (Tanaka et al. 2013) is not analyzed in this study.
<sup>c</sup> Ic-5 and Ic-6 shows identical genotype, since InDel2 (Tanaka et al. 2013) is not analyzed in this study.

A phylogenetic relationship of the three cytoplasm types, Ia, Ib and Ic, was demonstrated by the MJ network in which four wild species of *Cucumis* were included as outgroups (Fig. 3). A total of 15 subtypes detected in *C. melo* formed three distinct groups with no reticulation among them (i.e., no intermediates between the three groups). Therefore, the three cytoplasm types were distinctly differentiated from each other. In the MJ network, the Ia and Ib types were allocated in the same lineage, with the Ia type positioned closer to the outgroups, suggesting that it was cultivated earlier than the Ib type and that the Ib type evolved later. On the other hand, the Ic type evolved from wild species of *Cucumis* in another lineage. Interestingly, two subtypes, Ic-5 and Ic-6, were found in Ghana and Senegal, in the west of the prime meridian, while the other four subtypes of Ic were in the east. The tree topology, (Ic, (Ib,Ia)), was also confirmed in the NJ and ML trees (Supplemental Fig. 1).

**Fig. 3.**
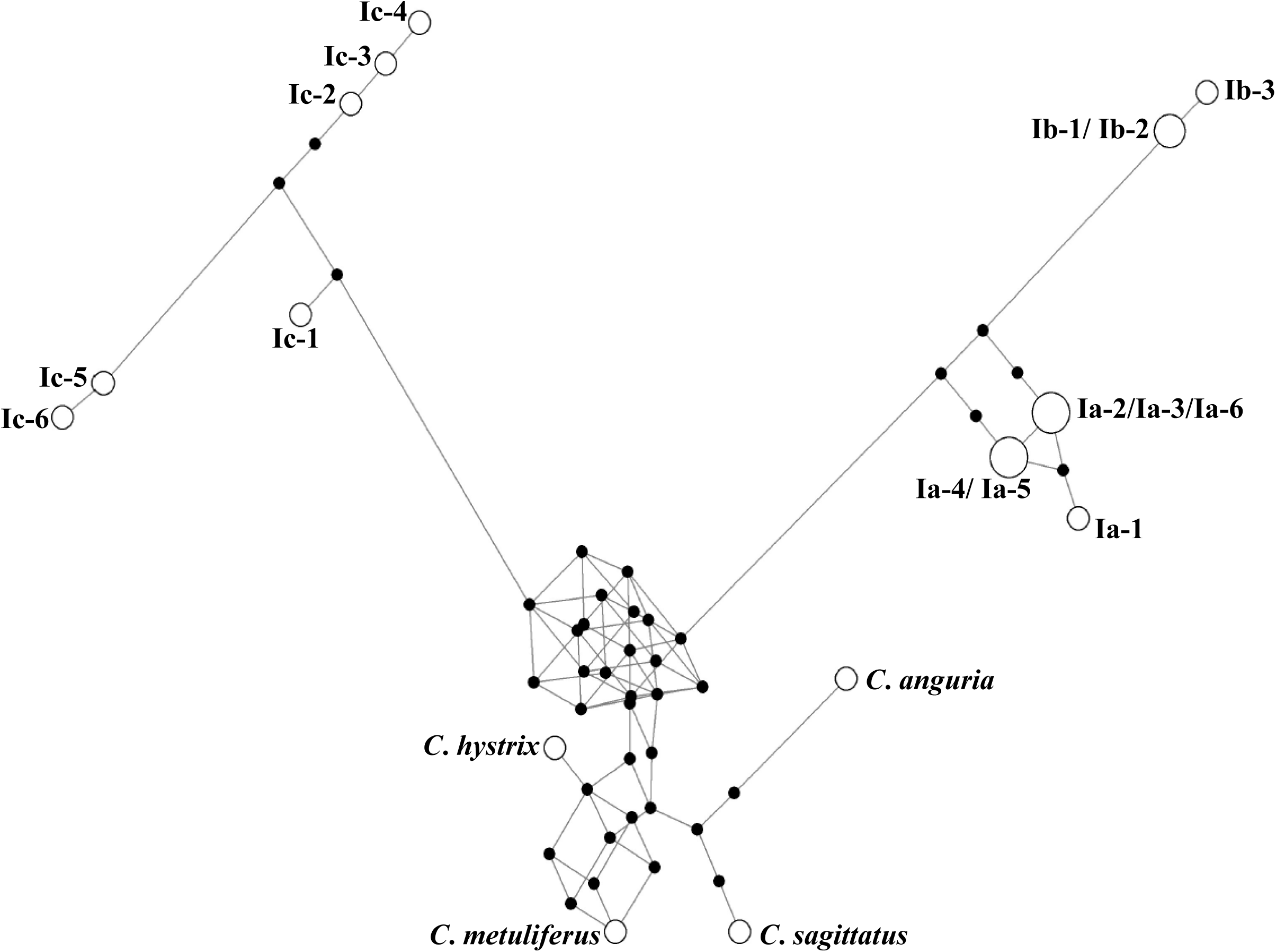
MJ networks (ε = 10) of 19 types of chloroplast genome in *Cucumis*. White open circles indicate each chloroplast type. The sequence polymorphisms on the link (SNP1-35, InDel 1–5) refer to mutated nucleotides. Sequence types indicated by black solid circles are added for growing network as median vectors.

### Nuclear marker genotyping

Eighteen RAPD primers amplified 27 polymorphic marker bands of approximately 550– 2027 bp, with an average of 1.7 effective marker bands produced by each primer (Supplemental Table 3). PIC varied depending on the primer, and ranged from 0.044 to 0.282. The most polymorphic marker band was B84-600 (PIC = 0.282), followed by A41-930 (PIC = 0.257) and B32-900 (PIC = 0.252).

Gene diversity (*D*) ranged from 0.147 to 0.367 in 10 melon groups classified based on geographical origin and seed length type (Supplemental Table 5). The largest diversity (*D* = 0.367) was observed in the large seed-type of South Asia. For small seed-type melon, the largest diversity (*D* = 0.292) was observed also in South Asia. In both large and small seed types, the *D* values gradually decreased from India towards Central and West Asia and Europe on the west, and towards Southeast Asia and East Asia on the east.

LnP(D) values from the STRUCTURE analysis increased with *K* from 2 to 10, with an evident inflection at *K* = 2 (Fig. 4A). According to second-order statistics, to estimate the number of subpopulations, the optimal value of *K* = 2 was identified, at which Delta *K* (Evanno et al. 2005) showed a peak (Fig. 4B). These results indicate that the diverse melon accessions used in this study consisted of two populations, designated PopA and PopB (Fig. 4C). Accessions with an estimated membership of over 0.5 were assigned to the respective population. Subpopulation structuring under PopA and PopB was performed by repeating the simulation for each population, and 12 accessions with estimated memberships below 0.7 were assigned to the “admixed group” (Supplemental Table 1). PopA was divided into three subgroups (*K* = 3) and PopB into two subgroups (*K* = 2) (Supplemental Fig. 2AB). Consequently, the 212 accessions were divided into five subgroups designated as PopA1 (53 accessions), PopA2 (32 accessions), PopA3 (eight accessions), PopB1 (72 accessions), and PopB2 (35 accessions), except for the 12 accessions of the admixed group. Genetic divergence, estimated by genetic distance and *F_st_* value, was rather small between PopA1 and PopA2 as well as PopB1 and PoB2, while PopA3 was highly divergent from the other populations (Supplemental Table 6).

**Fig. 4.**
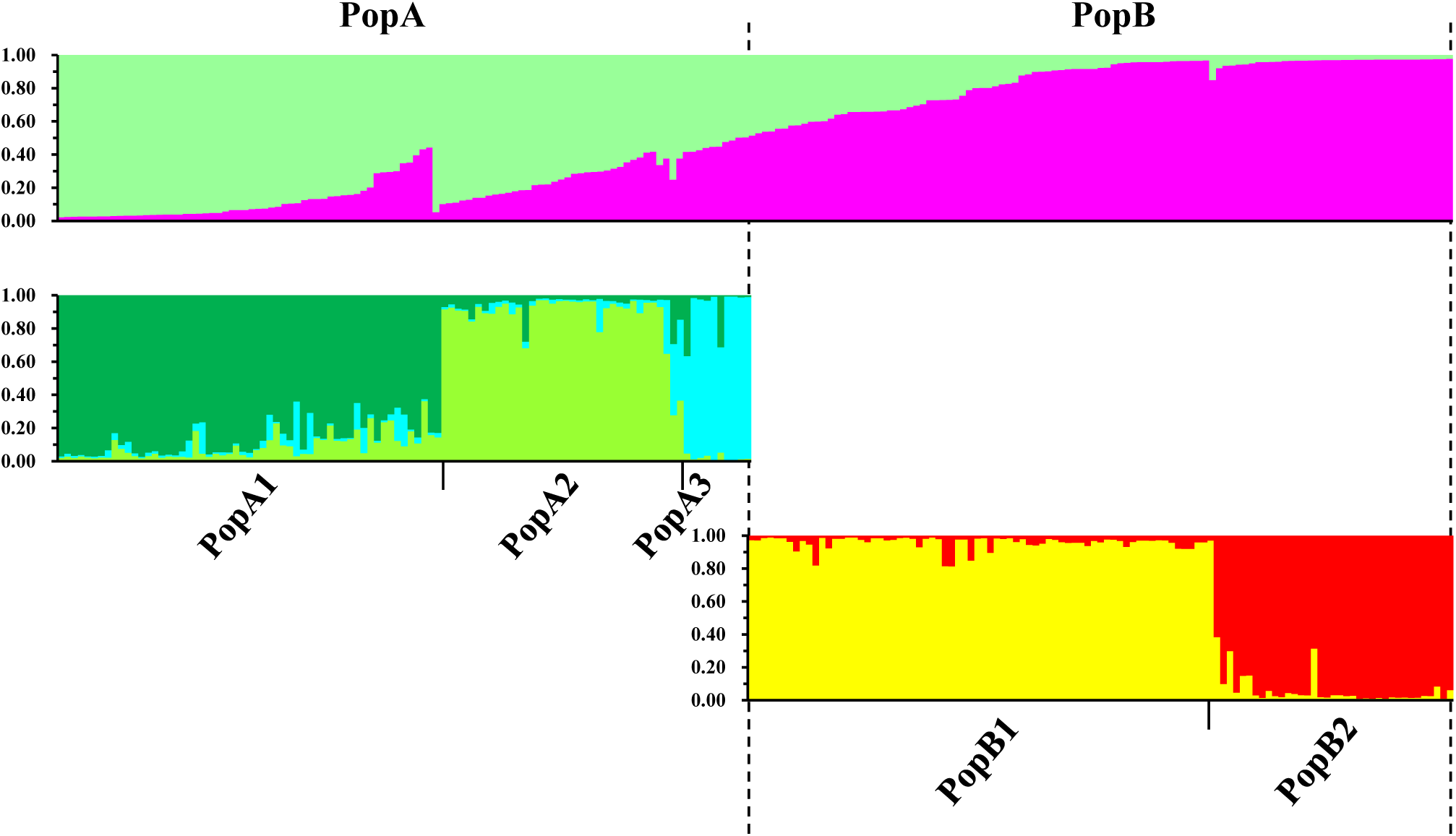
Classification of 212 melon accessions by STRUCTURE simulation based on RAPD and SSR polymorphisms. A) The ad hoc procedure described by Pritchard *et al*. (2000). B) The second order of statistics (Delta *K*) based on Evanno *et al*. (2005). C) Optimal population structure (*K* = 2) and subpopulation structure of PopA (*K* = 3) and PopB (*K* = 2). Each single vertical line represents a melon accession, and different colors represent different populations and subpopulations. The length of the colored segment illustrates the estimated proportion of each accession’s membership in the corresponding groups.

An unrooted UPGMA tree of 212 accessions was constructed, based on GD calculated after Apostol et al. (1993). It illustrated genetic relationships that closely approximated the STRUCTURE-based membership assignment for most accessions (Fig. 5A). For three subgroups, PopA1, PopA3, and PopB2, accessions of each subgroup formed the respective cluster(s), while accessions of PopA2 and PopB1 were classified into several clusters together with those of other subgroups. Close relationships were again demonstrated between PopA1 and PopA2, and between PopB1 and PopB2. A PCO was conducted to assess the adequacy of population subdivisions by STRUCTURE (Fig. 5B). Of the total variance among the 212 accessions, the first and second principal coordinates explained 43.77% and 15.23%, respectively. Plotting of the first two principal components showed separation of inferred subpopulations, which was highly consistent with STRUCTURE, and showed relationships of these subpopulations, which was consistent with the UPGMA tree. The relationships of all five subpopulations suggested by the UPGMA tree and PCO were also concordant with the GD and *F_ST_* (Supplemental Table 6).

**Fig. 5.**
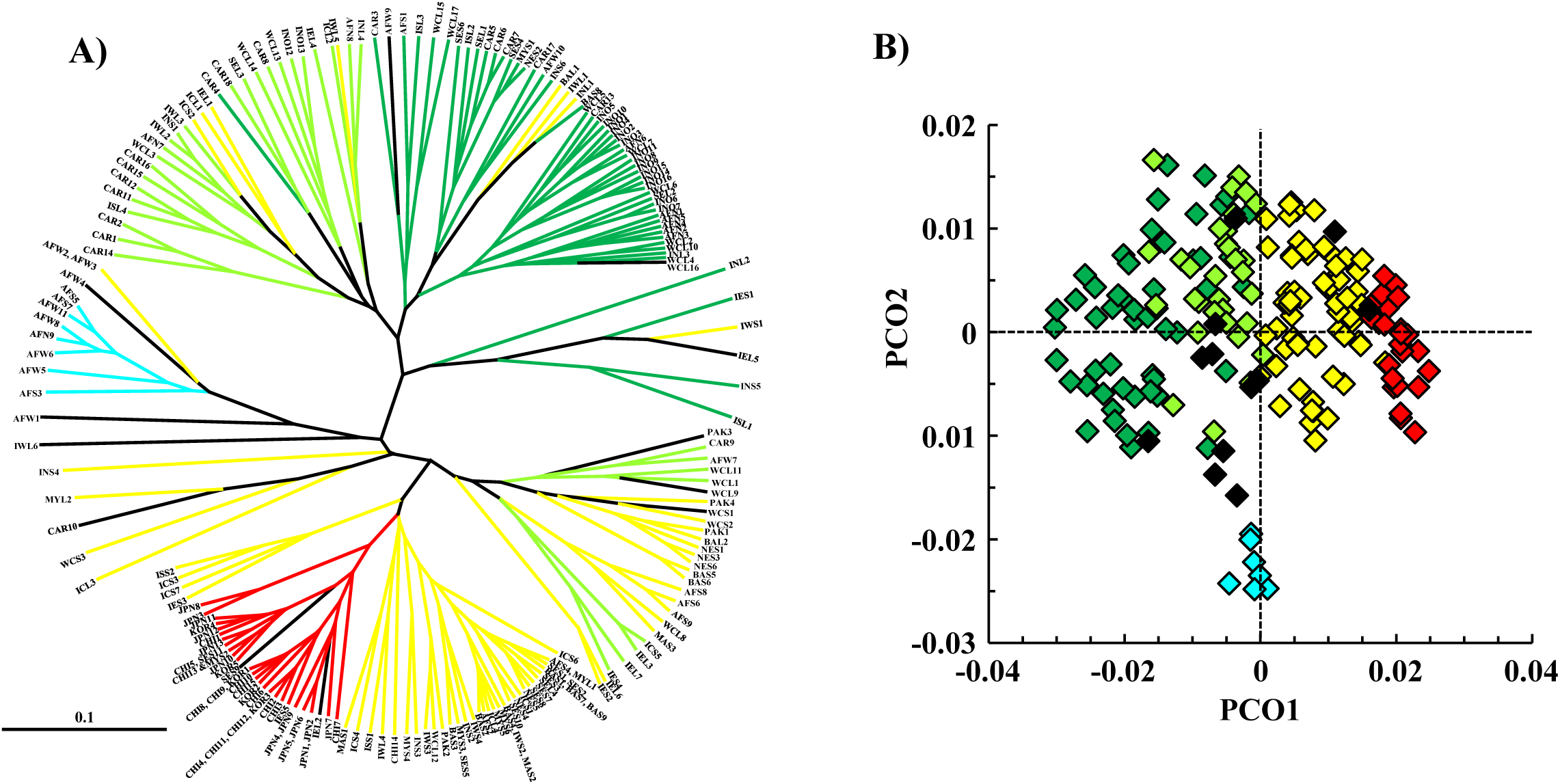
Genetic relationships of 212 melon accessions inferred from RAPD and SSR polymorphisms. A) UPGMA tree of 212 melon accessions. Colors indicate the STRUCTURE-based subpopulations: Green = PopA1, light green = Pop A2, light blue = Pop A3, yellow = PopB1, red = PopB2. B) Distribution of the STRUCTURE-based subpopulations on the first two principal co-ordinates. Colors indicate subpopulations as mentioned above.

### Classification based on seed length, chloroplast genome and nuclear genome

A total of 212 accessions were consequently classified into several groups as follows: two groups by seed length (Fig. 1), three groups by chloroplast DNA polymorphism (Fig. 3), and five populations by nuclear DNA polymorphism (Fig. 4C). If these three classifications are genetically independent, then at most 30 types, except for admixture types, are possible by combining three classifications. However, 200 of the 212 accessions (excluding 12 accessions of admixed groups) were assigned to only 17 types, of which three types consisted of one accession, suggesting non-random association between three classifications (Table 4). For simplicity, the results of the 200 accessions are indicated hereafter, unless otherwise stated.

**Table 4.** Association between three different classifications based on seed length and polymorphisms in chloroplast and nuclear genomes.

| Area / Seed size | Number of accessions | Ia<sup>a</sup> |  |  |  |  |  | Ib<sup>a</sup> |  |  |  |  |  | Ic<sup>a</sup> |  |  |  |  |  | Gene diversity |
| --- | --- | --- | --- | --- | --- | --- | --- | --- | --- | --- | --- | --- | --- | --- | --- | --- | --- | --- | --- | --- |
|  |  | A1 | A2 | A3 | B1 | B2 | AD | A1 | A2 | A3 | B1 | B2 | AD | A1 | A2 | A3 | B1 | B2 | AD |  |
| Europe, USA<sup>b</sup> |  |  |  |  |  |  |  |  |  |  |  |  |  |  |  |  |  |  |  | 0.308 |
| Large seed | 27 | - | - | - | - | - | - | 16 | 11 | - | - | - | (1) | - | - | - | - | - | - |  |
| Small seed | 1 | - | - | - | - | - | - | - | 1 | - | - | - | - | - | - | - | - | - | - |  |
| West and Central Asia |  |  |  |  |  |  |  |  |  |  |  |  |  |  |  |  |  |  |  | 0.345 |
| Large seed | 20 | - | 2 | - | 1 | - | - | 13 | 3 | - | 1 | - | (2) | - | - | - | - | - | - |  |
| Small seed | 2 | - | - | - | 2 | - | (1) | - | - | - | - | - | - | - | - | - | - | - | - |  |
| East Asia |  |  |  |  |  |  |  |  |  |  |  |  |  |  |  |  |  |  |  | 0.171 |
| Large seed | 0 | - | - | - | - | - | - | - | - | - | - | - | - | - | - | - | - | - | - |  |
| Small seed | 33 | - | - | - | 1 | 31 | - | - | - | - | - | 1 | - | - | - | - | - | - | - |  |
| South and Southeast Asia |  |  |  |  |  |  |  |  |  |  |  |  |  |  |  |  |  |  |  | 0.338 |
| Large seed | 29 | 2 | 3 | - | 8 | - | - | 8 | 4 | - | 4 | - | (2) | - | - | - | - | - | - |  |
| Small seed | 62 | 3 | 5 | - | 46 | 3 | (3) | 3 | - | - | 2 | - | - | - | - | - | - | - | - |  |
| Africa |  |  |  |  |  |  |  |  |  |  |  |  |  |  |  |  |  |  |  | 0.312 |
| Large seed | 11 | 1 | - | - | - | - | - | 7 | 3 | - | - | - | - | - | - | - | - | - | - |  |
| Small seed | 15 | - | - | 1 | 5 | - | - | - | - | - | - | - | - | - | - | 7 | 2 | - | (3) |  |
| Total | 200 | 6 | 10 | 1 | 63 | 34 | (4) | 47 | 22 | - | 7 | 1 | (5) | - | - | 7 | 2 | - | (3) |  |

<sup>a</sup> Ia, Ib and Ic indicate cytoplasm type. A1, A2, A3, B1 and B2 indicate model-based subpopulation, and AD indicates admixture type. No. of admixture accessions indicated in parentheses.
<sup>b</sup> East Asian *Cantalupensis* and *Inodorus* are included.
<sup>c</sup> Five types representing Europe/US large seed-type, East Asian small seed-type, and Ic type of Africa are highlighted in green, pink, and blue, respectively.

Among 61 accessions from Europe/US and East Asia, 59 (96.7%) were classified into four contrasting types: [large seed, Ib, PopA1 or A2] and [small seed, Ia, PopB1 or B2], highlighted in Table 4. Intermediate or recombinant types which could be derived from hybridization between those four types were less frequent. These results show distinct genetic differentiation between the Europe/US large seed type and the East Asian small seed type. Although many Europe/US and West and Central Asian melon were classified into two major types [large seed, Ib, PopA1 or A2], the subtype classified differed among horticultural groups, with European Cantalupensis as [Ib-1, PopA1 or A2] and British Cantalupensis and Chinese Hami melon of Inodorus as [Ib-2, PopA1] (Supplemental Table 1). Within the USA, Honeydew melon of Inodorus was characterized by [Ib-3, PopA1] and Cantalupensis mostly by [Ib-2 or Ib-3, PopA2].

Four contrasting types, [large seed, Ib, PopA1 or A2] and [small seed, Ia, PopB1 or B2], were also commonly found in South and Southeast Asia (Table 4, Supplemental Table 1). However, the percentage of these four types was 67.0% (61 of 91 accessions), which was lower than that in Europe/US and East Asia. Instead, recombinant types between these four contrasting types were rather frequent (33.0%, 30/91), and accessions of large and small seed types consisted of the respective six types. Among the 30 recombinants, five (16.7%) were classified as [large seed, Ia, PopA1 or A2], having the characteristics of both the large seed-type (PopA1 or A2) and the small seed-type (Ia). In addition, the frequencies of recombinant types of accessions were 17 among 29 accessions (58.6%) in the large seed-type, and 13 among 62 accessions (21.0%) in the small seed-type. These results indicated that recombinant types were found more frequently in the large seed-type than in the small seed-type. Interestingly, of the 17 recombinant accessions of the large seed-type, 12 were classified as either Momordica (seven accessions) or Flexuosus (five).

The unique Ic cytoplasm, which was not found outside Africa, was found frequently in Africa (41.4%, 12 of 29 accessions including admixed groups), though the number of accessions examined was limited (Table 4). The unique type [small seed, Ic, PopA3] accounted for 24.1% of melon accessions from Africa and 38.9% of African small seed type. In addition, three accessions of admixed groups proved to be [small seed, Ic, admixture PopA1/A3] and commonly shared the membership of PopA3. Four contrasting types commonly found worldwide, [large seed, Ib, PopA1 or A2] and [small seed, Ia, PopB1 or B2], were also detected (10 and five accessions, respectively). These results indicated that five types of melon, including the unique Ic type, are cultivated or grow in Africa, and that they are differentiated as a set of cytoplasm and nuclear genomes. More interestingly, these five types were found allopatrically: eight of 10 accessions of [large seed, Ib, PopA1 or A2] were in Northern Africa, all accessions of [small seed, Ia, PopB1] were in Southern Africa, and five of seven accessions of [small seed, Ic, PopA3] were in Western Africa.

## Discussion

Genetic relationships and divergence among geographical populations and horticultural varieties have been studied in melon, and it is widely accepted that the Europe/US melon (subsp. *melo*) and East Asian melon (subsp. *agrestis*) are genetically divergent in the nuclear genome (Silberstein et al. 1999; Monforte et al. 2003; Deleu et al. 2009; Esteras et al. 2009; Blanca et al. 2011; Sabato et al. 2015; Zhao et al. 2019; Liu et al. 2020; Wang et al. 2021; Shigita et al. 2023). The Europe/US melon and East Asian melon were separated by sequence polymorphisms in the chloroplast genome (Tanaka et al. 2013; Zhu et al. 2016; Endl et al. 2018; Zhao et al. 2019). In the present study we analyzed sequence polymorphism of the chloroplast genome by a PCR-based method, and identified three cytoplasm types, Ia, Ib, and Ic (Table 3), as previously reported (Tanaka et al. 2013). Zhao et al. (2019) analyzed genomic variation by whole genome resequencing, using 1,175 melon accessions, and classified them into three clades, *agrestis* clade, *melo* clade, and African clade, based on chloroplast SNPs. These three clades corresponded well to the three cytoplasm types reported by Tanaka et al. (2013). Zhu et al. (2016) classified seven melon accessions into two groups by the analysis of 63 SNPs in the chloroplast genome. Of these SNPs, six were identical to the SNP markers analyzed in this study (SNPs 2, 8, 18, 19, 25 and 29), by which the three cytoplasm types could be identified (Table 3). Since the Ic type accession of Africa was not included in the study by Zhu et al. (2016), seven accessions were separated into two clades, likely corresponding to Ia and Ib. Therefore, it was confirmed that the 12 markers (Table 1) developed in this study are applicable to the classification of chloroplast genome type, and that the set of markers, Cmcp1-dCAPS, Cmcp6-dCAPS, and Cmcp11-CAPS2, is sufficient to classify the three major cytoplasm types, Ia, Ib, and Ic.

Based on the analysis of seed size and molecular polymorphisms of nuclear and chloroplast genomes, using melon accessions covering wide geographical areas of the world, it was examined whether seed size and nuclear and chloroplast genomes diversified with close association between them. For the Europe/US melon and East Asian melon, all accessions of Europe/US were classified as [large seed, Ib, PopA1 or A2] except for one Japanese breeding line, while 32 of 33 accessions of East Asia were classified as [small seed, Ia, PopB1 or B2] (Table 4). In addition, recombinant types between the four contrasting types were rarely found in these areas, indicating nearly perfect divergence in both nuclear and chloroplast genomes between two peripheral areas of melon distribution. These results support independent origin of subsp. *melo* of Europe/US and subsp. *agrestis* of East Asia (Zhao et al. 2019). A large genetic diversity of melon proved to be partly ascribable to its polyphyletic origin.

In contrast, less attention has been paid to cultivated melon in South Asia and only little is known about their maternal lineage, despite India being known as the secondary center of diversity of cultivated melon (Robinson and Decker-Walters 1997; Akashi et al. 2002; Tanaka et al. 2007) and also claimed as the place of melon domestication (Sebastian et al. 2010; John et al. 2013; Endl et al. 2018; Zhao et al. 2019). In South and Southeast Asia, both large and small seed types are commonly found, and seeds length varied continuously from 3.9 mm to 13.2 mm (Fig. 1). More importantly, accessions with seed lengths from 8 mm to 10 mm, flanking the boundary between large and small seed-types, accounted for 30.2%, which was higher than those in Europe/US (14.7%) and East Asia (6.0%). The analysis of 1,998 Indian accessions of USDA-ARS GRIN showed a similar result (Table 2), confirming the result shown in Fig. 1. In this context, Momordica and Flexuosus included 15 accessions of large seed-type and seven accessions of small seed-type (Supplemental Table 1), and their seed lengths showed continuous variation from 5.7 mm to 13.2 mm across the boundary between large and small seed types. These two groups enrich the diversity in seed lengths, and make it difficult to classify South and Southeast Asian melon accessions into subsp. *melo* and subsp. *agrestis* merely by seed length.

The large genetic variation in South and Southeast Asia was confirmed by the presence of four types, [large seed, Ib, PopA1 or A2] of subsp. *melo* and [small seed, Ia, PopB1 or B2] of subsp. *agrestis*, and of the recombinants among them (Table 4). Except for South and Southeast Asia and Africa, [large seed, Ib, PopA1 or A2] comprised 91.5% of the large seed-type, and [small seed, Ia, PopB1 or B2] accounted for 94.4% of the small seed-type. In contrast, in South and Southeast Asia, their proportions were low, being 41.4% in the large seed-type and 79.0% in the small seed-type, showing the abundance of recombinant/intermediate types. As a result, no distinct genetic differentiation was detected in South and Southeast Asia. From these results, it is indicated that South and Southeast Asia, especially India, is rich in genetic diversity of seed length and nuclear and chloroplast genomes (Fig. 1, Supplemental Table 5), as reported previously (Akashi et al. 2002; Dhillon et al. 2007; Tanaka et al. 2007; Gonzalo et al. 2019; Shigita et al. 2023). Genetic variation is expected to be large simply by the coexistence of different types of melon, including Momordica, even if they are reproductively isolated from each other. However, genetic interchange through spontaneous hybridization between subsp. *melo* and subsp. *agrestis* seems to be common in India, especially in the large seed type, showing that polymorphisms of seed length and nuclear and chloroplast genomes are not closely associated, unlike the Europe/US subsp. *melo* and East Asian subsp. *agrestis* (Table 4). The presence of various kinds of recombinant/intermediate types made their botanical classification difficult, and supported the practical classification into horticultural groups proposed by Pitrat (2016). The Indian origin of melon with Ia and Ib cytoplasms was also affirmed by such an intermixed structure of genetic variation.

Momordica is mainly grown in India and Southeast Asia, and is characterized by unique characteristics such as mealy flesh, very thin exocarp splitting at maturity, and so on (Dhillon et al. 2007; Pitrat 2016). Fujishita (2008) and Tanaka et al. (2015) analyzed the length of ancient melon seed remains excavated from various sites in Japan, and found a historical transition in seed length. All but one of the 800 seed remains were shorter than 8.1 mm in length before the fourth century (5.7 mm on average), suggesting the utilization of Agrestis, and possibly Conomon and Makuwa. In contrast, from the fourth century to the eleventh, seed remains measuring over 8.1 mm became common (44.5%) and those shorter than 6.1 mm became rare (5.6%) (7.8 mm on average), suggesting a replacement with Momordica introduced from the continent. Thereafter, seed remains from 6.1 mm to 8.0 mm became predominant (77.1%), suggesting a replacement with Conomon and Makuwa introduced from the continent. Reflecting such a history, Momordica (locally called Shimauri or Babagoroshi) is still cultivated on Hachijojima Island in Japan, a remote island located 287 km south of Tokyo, and shows the same characteristics as Indian Momordica. These results indicate the spread of Momordica to Japan by the fourth century and its wide distribution in Asia in the past, though it is now limited to South and Southeast Asia. Of eight accessions of Indian Momordica examined in the present study, four (50.0%) proved to be the recombinant type. Both Ia and Ib types were detected. These results, together with seed length variation, suggest that Momordica might have originated from the reciprocal hybrids between [large seed, Ib, PopA1/A2] and [small seed, Ia, PopB1/B2] types. Flexuosus might have been similarly established.

Africa is considered to be one domestication center of melon (Zohary and Hopf 1988; Kirkbride 1993; Pitrat 2008; Endl et al. 2018; Zhao et al. 2019). As shown in Table 4 and Supplemental Table 1, allopatric distribution of three groups of melon was confirmed in Africa; [large seed, Ib, PopA1 or A2], [small seed, Ia, PopB1], and [small seed, Ic, PopA3], distributed mainly in Northern, Southern, and Western Africa, respectively. Tanaka et al. (2013) identified melon accessions with Ic cytoplasm only in the African continent, mainly in Western Africa but with a few in Southern Africa. Although seven additional accessions from Africa were examined in this study, all were of Ia or Ib type, and thus, so far, Ic type appears to be endemic to Western and Southern Africa. These results indicate distinct genetic differentiation among three groups of melon, subsp. *melo* and the Ia and Ic types of subsp. *agrestis*, indicating nearly perfect divergence in both nuclear and chloroplast genomes. The MJ network, NJ tree, and ML tree based on SNPs in the chloroplast genome place the Ic type apart from the Ia and Ib types (Fig. 3, Supplemental Fig. 1). Endl et al. (2018) located Tibish, a unique type of vegetable melon endemic to Sudan, in the most basal clade within *C*. *melo*, consisting exclusively of African accessions, and described this clade as subsp. *meloides*, which was considered the wild progenitor of Tibish and Fadasi. Among seven accessions of [small seed, Ic, PopA3] and three accessions of [small seed, Ic, admixture PopA1/A3], five were cultivated and five were wild or feral (Supplemental Table 1). A Ghanaian accession, PI 185111, classified as subsp. *meloides* by Endl et al. (2018), was identified as a [small seed, Ic, admixture PopA1/A3] type (Supplemental Table 1), suggesting a close relationship between subsp. *meloides* and Ic type accessions of subsp. *agrestis*. The Ic type melon was considered to be domesticated from the Ic type wild melon subsp. *meloides* in Africa. Although Ic type melons such as Tibish and Fadasi are grown only in limited areas of Africa, they have a unique nuclear genome of PopA3 and thus could be of considerable potential value as unique genetic resources for breeding. Further studies using more accessions from Africa should be carried out.

## Author Contribution Statement

KT and KK conceived the project; KK provided materials; KT, TPD, PTPN, and MT performed the experiments; KT analyzed the data, prepared figures, and wrote a draft of the manuscript; and GS, YM, HN and RI provided advice on the experimental implementation and helped draft the manuscript.

## Acknowledgments

This work was partly supported by JSPS KAKENHI Grant Numbers 26257409 and 24K08891.

