## Supplemental figures for "Polyphyletic domestication and inter-lineage hybridization magnified genetic diversity of cultivated melon, *Cucumis melo* L"

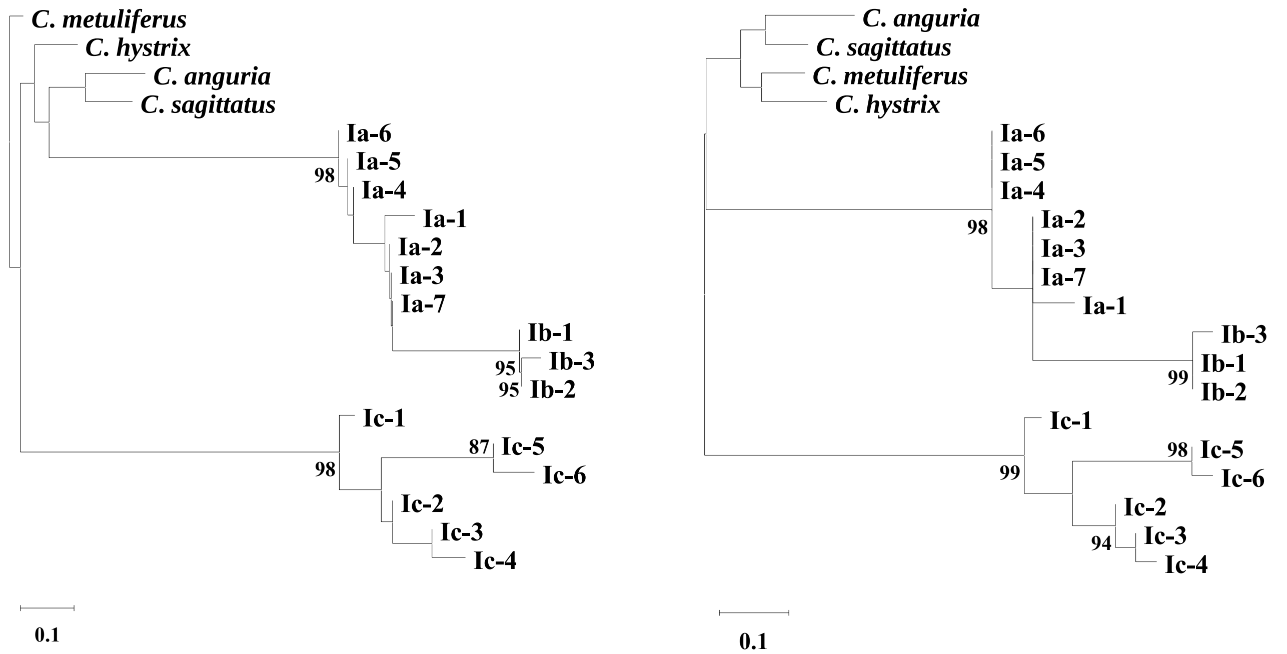


Supplemental Fig. 1. Neighbor-joining tree (left) and the maximum likelihood tree (right) for 19 cytoplasm types in *Cucumis*, based on sequence polymorphism in the chloroplast genome. Bootstrap values over 80% for 1000 replicates are shown beside the branch.

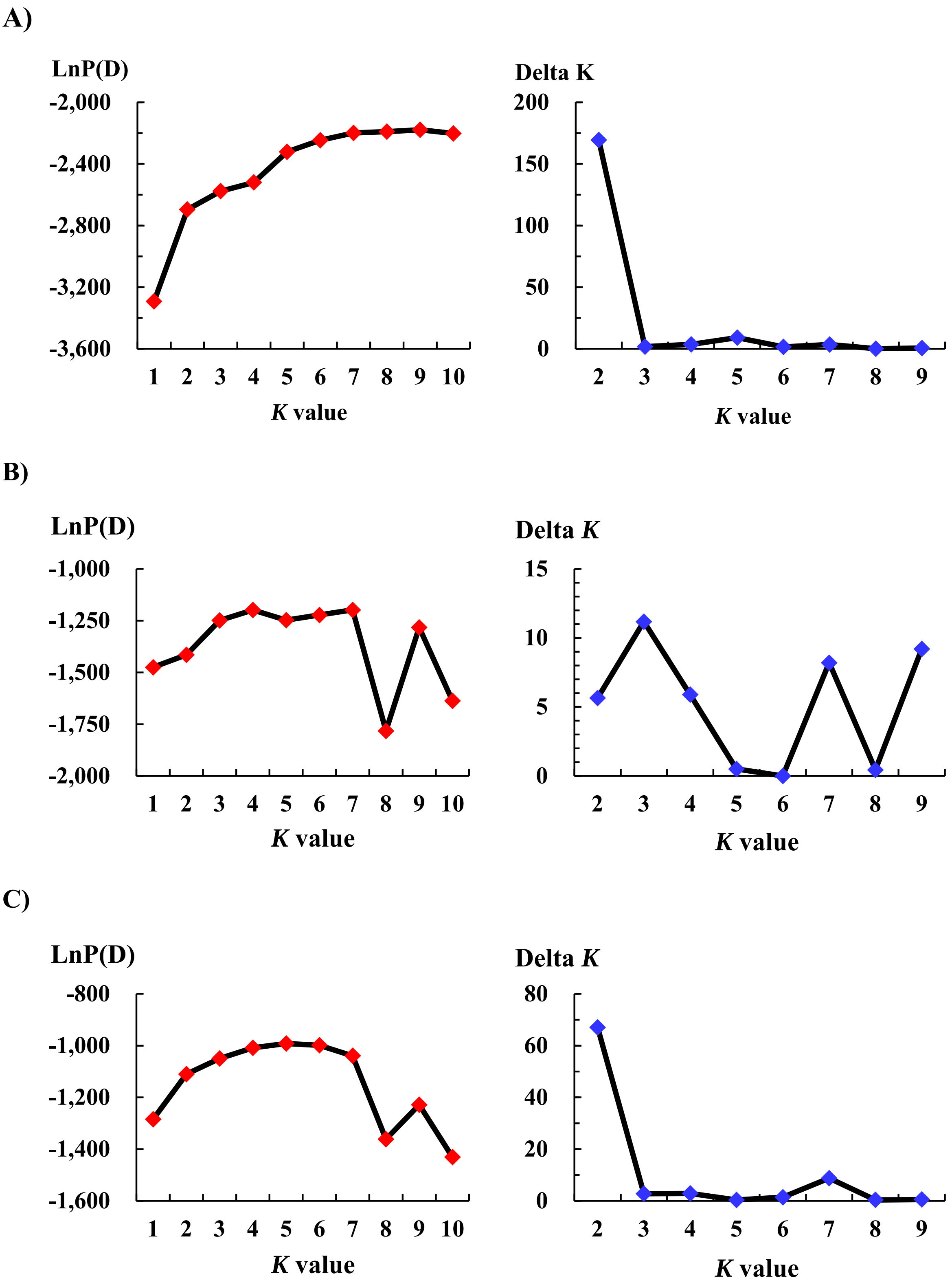


Supplemental Fig. 2. Determinations of value of *K* for substructuring in two model-based populations. A) Optimal *K* identified by LnP(D) (left side) and delta *K* (right side) in 212 melon accessions. B) Optimal *K* identified by LnP(D) (left side) and delta *K* (right side) in 103 melon accessions in PopA. C) Optimal *K* identified by LnP(D) (left side) and delta *K* (right side) in 109 melon accessions in PopB.

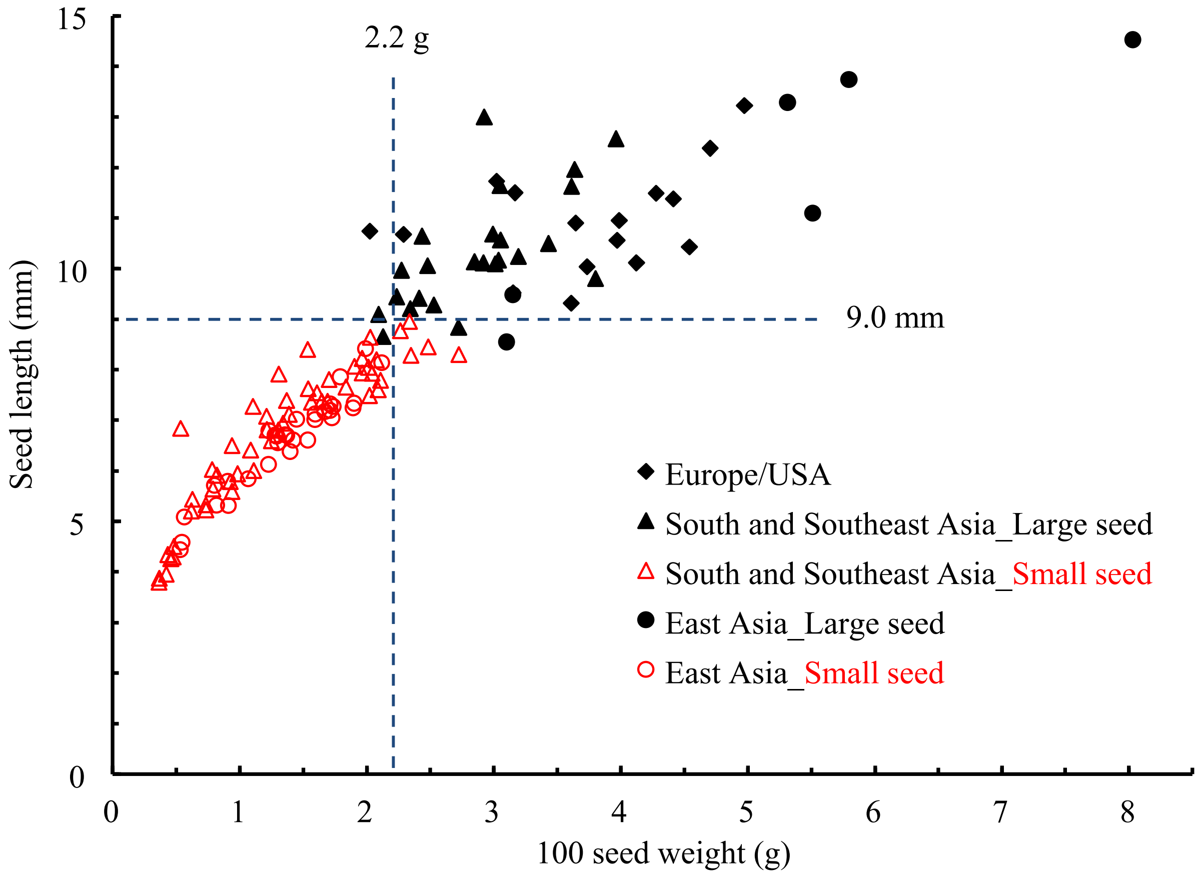


Supplemental Fig. 3. Relationship between weight and length of seeds in 130 accessions of melon.
