## Supplemental tables for "Polyphyletic domestication and inter-lineage hybridization magnified genetic diversity of cultivated melon, *Cucumis melo* L"

Supplemental Table 1. List of melon accessions analysed in this study

| Cultivar name / Accession number | Seed source [c] | Geographical group | Horticultural group | Cultivar group / Country | Seed type [d] | Chloroplast genome type | Cluster No. (UPGMA) | Membership (STRUCTURE) | 100 seed weight (g) |
|---|---|---|---|---|---|---|---|---|---|
| Melon Cantalupo di Charentais | 1 | Europe/US | Cantalupensis | Europian cantaloup | L | Ib-1 | IIc | Pop A2 | 3.97 |
| Cantaloure de Bellegrade [a] | 1 | Europe/US | Cantalupensis | Europian cantaloup | L | Ib-1 | IIc | Pop A2 | 3.99 |
| Ogen (780045) [a] | 1 | Europe/US | Cantalupensis | Europian cantaloup | L | Ib-1 | Ic | Pop A1 | 4.12 |
| Barnett Hill Favourite | 1 | Europe/US | Cantalupensis | England greenhouse type | L | Ib-2 | IIb | Pop A1 | – |
| Earl's Favorite | 1 | Europe/US | Cantalupensis | England greenhouse type | L | Ib-2 | Ib | Pop A1 | 3.61 |
| Blenheim Orange | 1 | Europe/US | Cantalupensis | England greenhouse type | L | Ib-2 | Ib | Pop A1 | 3.74 |
| British Queen | 1 | Europe/US | Cantalupensis | England greenhouse type | L | Ib-2 | Ib | Pop A1 | 3.15 |
| Hero of Lockinge | 1 | Europe/US | Cantalupensis | England greenhouse type | L | Ib-2 | IIb | Pop A2 | 4.54 |
| Homegarden | 1 | Europe/US | Cantalupensis | American field type | L | Ib-3 | VII | Pop A2 | 3.64 |
| Georgia 47 | 1 | Europe/US | Cantalupensis | American field type | L | Ib-2 | V | Pop A2/A3 | 2.03 |
| Rocky Ford | 1 | Europe/US | Cantalupensis | American field type | L | Ib-1 | IIc | Pop A2 | 4.28 |
| ♯58-21 | 1 | Europe/US | Cantalupensis | American field type | L | Ib-3 | IIc | Pop A2 | 4.41 |
| SC108 (C-108) | 1 | Europe/US | Cantalupensis | American field type | L | Ib-2 | Ia | Pop A1 | 3.02 |
| Rio Gold | 1 | Europe/US | Cantalupensis | American field type | L | Ib-2 | IIc | Pop A2 | – |
| Hale's Best | 1 | Europe/US | Cantalupensis | American field type | L | Ib-2 | IIc | Pop A2 | – |
| Spicy | 1 | Europe/US | Cantalupensis | American field type | L | Ib-3 | IIc | Pop A2 | – |
| Kurume 2 | 1 | Europe/US | Cantalupensis | Japan breeding line | L | Ib-2 | Ib | Pop A1 | 3.15 |
| Melon Chuukanbohon Nou 1 | 1 | Europe/US | Cantalupensis | Japan breeding line | S | Ib-2 | IIb | Pop A2 | 3.10 |
| Honeydew | 1 | Europe/US | Inodorus | Honeydew | L | Ib-3 | Ia | Pop A1 | 3.17 |
| 600011 | 1 | Europe/US | Inodorus | Haneydew | L | Ib-3 | Ia | Pop A1 | 2.29 |
| 610002 | 1 | Europe/US | Inodorus | Haneydew | L | Ib-3 | Ia | Pop A1 | 4.97 |
| 650013 | 1 | Europe/US | Inodorus | Haneydew | L | Ib-3 | Ia | Pop A1 | 4.71 |
| Chinese Honeydew | 1 | Europe/US | Inodorus | Chinese Honeydew | L | Ib-3 | Ia | Pop A1 | 5.31 |
| Carosello Scopatizzo Barese | 1 | Europe/US | Inodorus | Spain | L | Ib-2 | Ia | Pop A1 | – |
| Spain Noboru 3 | 1 | Europe/US | Inodorus | Spain | L | Ib-3 | IIb | Pop A2 | – |
| Tendral [b] | 1 | Europe/US | Inodorus | Spain | L | Ib-3 | IIb | Pop A2 | – |
| Kokand | 1 | Europe/US | Inodorus | Russia | L | Ib-3 | Ia | Pop A1 | – |
| Mirzuchulskaja | 1 | Europe/US | Inodorus | Russia | L | Ib-3 | Ia | Pop A1 | – |
| Ak-Urug | 1 | Europe/US | Inodorus | Russia | L | Ib-3 | Ia | Pop A1 | – |
| Hami-gua 6 | 1 | West and Central Asia | Inodorus | Chinese Hami melon | L | Ib-2 | Ia | Pop A1 | 8.03 |
| Hami-gua | 1 | West and Central Asia | Inodorus | Chinese Hami melon | L | Ib-2 | Ia | Pop A1 | 5.51 |
| Hami-gua 2 | 1 | West and Central Asia | Inodorus | Chinese Hami melon | L | Ib-2 | Ia | Pop A1 | 5.80 |
| Hami-gua H | 1 | West and Central Asia | Inodorus | Chinese Hami melon | L | Ib-2 | Ia | Pop A1 | – |
| Hami-gua J | 1 | West and Central Asia | Inodorus | Chinese Hami melon | L | Ib-2 | Ia | Pop A1 | – |
| 940068 | 1 | West and Central Asia | Agrestis | Iran | S | Ia-3 | VII | Pop A2/A3 | – |
| 940069 | 1 | West and Central Asia | Agrestis | Iran | S | Ia-3 | VII | Pop B1 | – |
| Karasu melon | 1 | West and Central Asia | Unclassified landrace | Turkey | L | Ib-2 | VII | Pop A2 | – |
| PI 169379 | 2 | West and Central Asia | Unclassified landrace | Turkey | L | Ib-1 | Ia | Pop A1 | – |
| PI 172818 | 2 | West and Central Asia | Unclassified landrace | Turkey | L | Ia-3 | IIc | Pop A2 | – |
| PI 176928 | 2 | West and Central Asia | Unclassified landrace | Turkey | L | Ib-2 | Ia | Pop A1 | – |
| PI 181872 | 2 | West and Central Asia | Unclassified landrace | Syria | L | Ib-2 | Ia | Pop A1 | – |
| PI 534605 | 2 | West and Central Asia | Unclassified landrace | Syria | L | Ib-1 | Ia | Pop A1 | – |
| PI 181747 | 2 | West and Central Asia | Unclassified landrace | Lebanon | L | Ib-2 | Ia | Pop A1 | – |
| PI 369161 | 2 | West and Central Asia | Unclassified landrace | Lebanon | L | Ia-1 | VII | Pop B1 | – |
| PI 435286 | 2 | West and Central Asia | Unclassified landrace | Iraq | S | Ia-3 | V | Pop B1 | – |
| PI 435290 | 2 | West and Central Asia | Unclassified landrace | Iraq | L | Ib-2 | VII | Pop A1/A2 | – |
| PI 140814 | 2 | West and Central Asia | Unclassified landrace | Iran | L | Ib-3 | Ia | Pop A1 | – |
| PI 143231 | 2 | West and Central Asia | Unclassified landrace | Iran | L | Ib-3 | VII | Pop A2 | – |
| PI 230185 | 2 | West and Central Asia | Unclassified landrace | Iran | L | Ib-2 | VI-c | Pop B1 | – |
| PI 126054 | 2 | West and Central Asia | Unclassified landrace | Afghanistan | L | Ia-3 | IIb | Pop A2 | – |
| PI 125931 | 2 | West and Central Asia | Unclassified landrace | Afghanistan | L | Ib-3 | IIb | Pop A2 | – |
| PI 125961 | 2 | West and Central Asia | Unclassified landrace | Afghanistan | L | Ib-1 | Ic | Pop A1 | – |
| PI 126047 | 2 | West and Central Asia | Unclassified landrace | Afghanistan | L | Ib-2 | Ia | Pop A1/A3 | – |
| PI 127532 | 2 | West and Central Asia | Unclassified landrace | Afghanistan | L | Ib-1 | Ic | Pop A1 | – |
| PI 614375 | 2 | South Asia | Flexuosus | India-west | L | Ia-1 | Ib | Pop A1 | – |
| PI 614543 | 2 | South Asia | Flexuosus | India-center | L | Ia-1 | IIc | Pop A2 | – |
| PI 614576 | 2 | South Asia | Flexuosus | India-west | L | Ia-1 | IIc | Pop A2 | 2.85 |
| NP20 | 3 | South Asia | Flexuosus | India-north | S | Ia-1 | IIc | Pop A2 | 1.54 |
| PI 614543 | 2 | South Asia | Flexuosus | India-center | L | Ia-1 | IIc | Pop B1 | 3.19 |
| NP12 | 3 | South Asia | Flexuosus | India-east | L | Ia-1 | IIc | Pop B1 | 2.72 |
| PI 182952 | 2 | South Asia | Momordica | India-west | L | Ib-1 | VI-c | Pop B1 | 3.05 |
| NP5 | 3 | South Asia | Momordica | India-north | S | Ia-1 | VI-c | Pop B1 | 1.11 |
| PI 116479 | 2 | South Asia | Momordica | India-north | L | Ia-1 | III | Pop A1 | 2.24 |
| NP14 | 3 | South Asia | Momordica | India-north | L | Ib-2 | Ib | Pop A1 | 2.48 |
| PI 614556 | 2 | South Asia | Momordica | India-center | L | Ib-1 | IIa | Pop A2 | 3.61 |
| PI 124096 | 2 | South Asia | Momordica | India-south | L | Ib-1 | VI-c | Pop B1 | 3.43 |
| PI 210077 | 2 | South Asia | Momordica | India-east | S | Ia-1 | VI-b | Pop B1/B2 | 2.48 |
| PI 124208 | 2 | South Asia | Momordica | India-east | L | Ia-1 | VII | Pop A2 | 3.96 |

| | | | | | | | | | |
|---|---|---|---|---|---|---|---|---|---|
| 940262 | 1 | South Asia | Momordica | Nepal | S | Ia-1 | VII | Pop B1 | 1.30 |
| 940263 | 1 | South Asia | Momordica | Nepal | S | Ib-1 | Ib | Pop A1 | 1.31 |
| 940275 | 1 | South Asia | Momordica | Bangladesh | S | Ia-3 | VI-c | Pop B1 | 0.63 |
| 790113 | 1 | South Asia | Momordica | Bangladesh | L | Ia-1 | Ib | Pop B1 | 2.13 |
| 760031 | 1 | South Asia | Momordica | Bangladesh | L | Ia-1 | VII | Pop B1 | 2.09 |
| 940099 | 1 | South Asia | Agrestis | Pakistan | S | Ia-4 | VII | Pop B1 | – |
| 940101 | 1 | South Asia | Agrestis | Pakistan | S | Ia-4 | VI-c | Pop B1 | – |
| 940102 | 1 | South Asia | Agrestis | Pakistan | S | Ia-5 | VII | Pop A1/A2 | – |
| 940103 | 1 | South Asia | Agrestis | Pakistan | S | Ia-4 | VII | Pop B1 | – |
| PI 164796 | 2 | South Asia | Agrestis | India-west | S | Ia-3 | III | Pop B1 | 0.62 |
| PI 614433 | 2 | South Asia | Agrestis | India-west | S | Ia-3 | VI-c | Pop B1 | 0.43 |
| NP10 | 3 | South Asia | Agrestis | India-north | S | Ia-1 | VI-c | Pop B1 | 0.73 |
| PI 614519 | 2 | South Asia | Agrestis | India-center | S | Ia-6 | VI-c | Pop B1 | 0.94 |
| PI 614549 | 2 | South Asia | Agrestis | India-center | S | Ia-1 | IIc | Pop B1 | 1.56 |
| PI 536481 | 2 | South Asia | Agrestis | Maldives | S | Ia-3 | VI-c | Pop B1 | – |
| 820003 | 1 | South Asia | Agrestis | Nepal | S | Ia-3 | VII | Pop B1 | 0.42 |
| 940097 | 1 | South Asia | Agrestis | Nepal | S | Ia-3 | VI-c | Pop B1 | 0.47 |
| 940109 | 1 | South Asia | Agrestis | Nepal | S | Ia-3 | VI-c | Pop B1 | 0.46 |
| 770134 | 1 | South Asia | Agrestis | Bangladesh | S | Ia-4 | VI-c | Pop B1 | 0.37 |
| 770135 | 1 | South Asia | Agrestis | Bangladesh | S | Ia-4 | VI-c | Pop B1 | 0.36 |
| 790112 | 1 | South Asia | Agrestis | Bangladesh | S | Ia-3 | VI-c | Pop B1 | 0.48 |
| PI 116738 | 2 | South Asia | Unclassified landrace | India-west | S | Ib-2 | VI-c | Pop B1 | 1.69 |
| PI 614262 | 2 | South Asia | Unclassified landrace | India-west | S | Ia-3 | VI-c | Pop B1 | 1.65 |
| PI 116666 | 2 | South Asia | Unclassified landrace | India-west | L | Ia-1 | IIa | Pop B1 | 2.34 |
| PI 164825 | 2 | South Asia | Unclassified landrace | India-west | L | Ib-1 | IV | Pop A1/A3 | 3.04 |
| PI 175109 | 2 | South Asia | Unclassified landrace | India-north | S | Ia-1 | V | Pop B1 | 2.01 |
| PI 179666 | 2 | South Asia | Unclassified landrace | India-north | S | Ib-1 | III | Pop A1 | 1.84 |
| NP2 | 3 | South Asia | Unclassified landrace | India-north | L | Ib-1 | Ib | Pop A1 | 3.01 |
| PI 124109 | 2 | South Asia | Unclassified landrace | India-north | L | Ib-1 | Ia | Pop A1 | 2.44 |
| PI 165508 | 2 | South Asia | Unclassified landrace | India-north | S | Ia-1 | IIa | Pop A2 | 1.96 |
| PI 614542 | 2 | South Asia | Unclassified landrace | India-center | S | Ia-1 | VI-a | Pop B1 | 2.02 |
| PI 614561 | 2 | South Asia | Unclassified landrace | India-center | S | Ia-2 | VI-c | Pop B1 | 2.35 |
| PI 614566 | 2 | South Asia | Unclassified landrace | India-center | S | Ia-1 | VII | Pop A2 | 2.09 |
| PI 614567 | 2 | South Asia | Unclassified landrace | India-center | S | Ia-1 | VI-c | Pop B1 | – |
| PI 614568 | 2 | South Asia | Unclassified landrace | India-center | S | Ia-1 | VI-a | Pop B1 | 0.54 |
| PI 124435 | 2 | South Asia | Unclassified landrace | India-center | L | Ib-2 | V | Pop B1 | 3.64 |
| PI 614588 | 2 | South Asia | Unclassified landrace | India-center | S | Ia-2 | VI-c | Pop B1 | 1.34 |
| PI 164323 | 2 | South Asia | Unclassified landrace | India-south | S | Ia-1 | VI-a | Pop B1 | 2.26 |
| PI 164585 | 2 | South Asia | Unclassified landrace | India-south | L | Ib-2 | III | Pop A1 | – |
| PI 123684 | 2 | South Asia | Unclassified landrace | India-south | L | Ib-3 | Ib | Pop A1 | 2.92 |
| PI 124105 | 2 | South Asia | Unclassified landrace | India-south | L | Ib-3 | Ic | Pop A1 | 3.05 |
| PI 123501 | 2 | South Asia | Unclassified landrace | India-south | L | Ib-2 | IIc | Pop A2 | 2.93 |
| PI 124113 | 2 | South Asia | Unclassified landrace | India-east | S | Ia-1 | III | Pop A1 | 2.03 |
| PI 124112 | 2 | South Asia | Unclassified landrace | India-east | S | Ib-1 | VII | Pop B1 | 1.11 |
| PI 166125 | 2 | South Asia | Unclassified landrace | India-east | S | Ia-1 | VI-a | Pop B1 | 0.78 |
| PI 124111 | 2 | South Asia | Unclassified landrace | India-east | L | Ib-1 | IIa | Pop A2 | 2.99 |
| PI 124207 | 2 | South Asia | Unclassified landrace | India-east | L | Ib-2 | III | Pop A1/A2 | 2.27 |
| PI 124112 | 2 | South Asia | Unclassified landrace | India-east | L | Ib-1 | VII | Pop B1 | – |
| PI 210541 | 2 | South Asia | Unclassified landrace | India-east | S | Ia-1 | VII | Pop A2 | 2.04 |
| PI 210542 | 2 | South Asia | Unclassified landrace | India-east | S | Ia-1 | VI-b | Pop B2 | 0.94 |
| PI 210541 | 2 | South Asia | Unclassified landrace | India-east | S | Ia-1 | VII | Pop A2 | – |
| PI 536479 | 2 | South Asia | Unclassified landrace | Maldives | S | Ia-1 | VI-c | Pop B1 | – |
| PI 536480 | 2 | South Asia | Unclassified landrace | Maldives | S | Ia-1 | VII | Pop B1 | – |
| NP1 | 3 | South Asia | Unclassified landrace | Nepal | S | Ia-1 | VII | Pop B1 | 2.07 |
| 790079 | 1 | South Asia | Unclassified landrace | Bangladesh | S | Ia-1 | VII | Pop B1 | 1.90 |
| Joydebpul | 1 | South Asia | Unclassified landrace | Bangladesh | S | Ia-1 | VII | Pop B1 | 1.70 |
| C5-3 | 1 | South Asia | Unclassified landrace | Bangladesh | S | Ia-1 | VI-c | Pop B1 | 0.73 |
| 890096 | 1 | South Asia | Unclassified landrace | Bangladesh | S | Ia-1 | Ib | Pop B1 | 1.96 |
| 940269 | 1 | South Asia | Unclassified landrace | Bangladesh | S | Ia-3 | VI-c | Pop B1 | 0.82 |
| 910050 | 1 | Southeast Asia | Flexuosus | Indonesia | L | Ib-1 | Ib | Pop A1 | 2.41 |
| 940261 | 1 | Southeast Asia | Momordica | Myanmar | S | Ia-1 | Ib | Pop A1 | 2.73 |
| 650057 | 1 | Southeast Asia | Momordica | Myanmar | L | Ia-1 | VI-c | Pop B1 | 2.53 |
| My238 | 3 | Southeast Asia | Unclassified landrace | Myanmar | S | Ia-1 | VI-b | Pop B2 | 1.25 |
| PI 200814 | 2 | Southeast Asia | Unclassified landrace | Myanmar | S | Ia-1 | VI-c | Pop B1 | 0.98 |
| PI 200816 | 2 | Southeast Asia | Unclassified landrace | Myanmar | S | Ia-1 | VI-c | Pop B1 | 2.11 |
| PI 200817 | 2 | Southeast Asia | Unclassified landrace | Myanmar | S | Ia-1 | VI-c | Pop B1 | 0.79 |
| PI 200819 | 2 | Southeast Asia | Unclassified landrace | Myanmar | S | Ia-1 | VI-c | Pop B1 | 1.61 |
| PI 200813 | 2 | Southeast Asia | Unclassified landrace | Myanmar | L | Ia-1 | V | Pop B1 | 3.80 |
| 940300 | 1 | Southeast Asia | Unclassified landrace | Thailand | S | Ia-3 | VI-c | Pop B1 | 1.54 |
| 940301 | 1 | Southeast Asia | Unclassified landrace | Thailand | S | Ia-3 | VI-c | Pop B1 | 1.39 |
| 940303 | 1 | Southeast Asia | Unclassified landrace | Thailand | S | Ia-1 | VI-c | Pop B1 | 1.37 |
| 940320 | 1 | Southeast Asia | Unclassified landrace | Thailand | S | Ib-1 | Ib | Pop A1 | 2.34 |
| T3 | 3 | Southeast Asia | Unclassified landrace | Thailand | S | Ia-1 | VI-c | Pop B1 | 1.21 |

| | | | | | | | | | |
|---|---|---|---|---|---|---|---|---|---|
| 860314 | 1 | Southeast Asia | Unclassified landrace | Thailand | S | Ia-1 | Ib | Pop A1 | 1.09 |
| 940289 | 1 | Southeast Asia | Unclassified landrace | Malaysia | S | Ia-1 | VI-c | Pop B1 | 0.93 |
| Horimatsu | 1 | Southeast Asia | Unclassified landrace | Malaysia | L | Ib-2 | Ia | Pop A1 | – |
| B1 | 3 | Southeast Asia | Unclassified landrace | Indonesia | S | Ia-1 | VI-c | Pop B1 | 1.21 |
| B2 | 3 | Southeast Asia | Unclassified landrace | Indonesia | S | Ia-1 | VI-b | Pop B1/B2 | – |
| BL1 | 3 | Southeast Asia | Unclassified landrace | Indonesia | L | Ib-3 | IIb | Pop A2 | – |
| MO-1 | 1 | Southeast Asia | Unclassified landrace | Laos | S | Ia-1 | VI-c | Pop B1 | – |
| DGT | 3 | Southeast Asia | Unclassified landrace | Viet Nam | L | Ia-1 | VI-c | Pop B1 | – |
| DHK | 3 | Southeast Asia | Unclassified landrace | Viet Nam | S | Ia-1 | VI-b | Pop B2 | – |
| C32 | 3 | East Asia | Conomon | China | S | Ia-1 | VI-b | Pop B2 | 1.40 |
| 940182 | 1 | East Asia | Conomon | China | S | Ia-3 | VI-b | Pop B2 | – |
| P169-1 | 3 | East Asia | Conomon | China | S | Ia-1 | VI-b | Pop B2 | 1.73 |
| P171-1 | 3 | East Asia | Conomon | China | S | Ia-1 | VI-b | Pop B2 | 1.73 |
| C28 | 3 | East Asia | Makuwa | China | S | Ia-1 | VI-b | Pop B2 | 1.54 |
| Mi-tang-tin | 1 | East Asia | Makuwa | China | S | Ia-1 | VI-b | Pop B2 | 1.59 |
| 760007 | 1 | East Asia | Makuwa | China | S | Ia-3 | VI-b | Pop B2 | 1.59 |
| 910008 | 1 | East Asia | Makuwa | China | S | Ia-1 | VI-b | Pop B2 | 1.71 |
| 780143 | 1 | East Asia | Makuwa | China | S | Ia-1 | VI-b | Pop B2 | 1.30 |
| 910055 | 1 | East Asia | Makuwa | China | S | Ia-1 | VI-b | Pop B2 | 1.37 |
| 940178 | 1 | East Asia | Makuwa | China | S | Ia-1 | VI-b | Pop B2 | 1.67 |
| 940184 | 1 | East Asia | Makuwa | China | S | Ia-1 | VI-b | Pop B2 | 1.89 |
| Chi-87-12 | 1 | East Asia | Makuwa | China | S | Ia-1 | VI-b | Pop B2 | 0.90 |
| PI 136173 | 2 | East Asia | Makuwa | China | S | Ia-1 | VI-c | Pop B1 | 1.36 |
| PI 157070 | 2 | East Asia | Makuwa | China | S | Ia-1 | VI-b | Pop B2 | 1.07 |
| 630044 | 1 | East Asia | Makuwa | Korea | S | Ia-1 | VI-b | Pop B2 | 1.23 |
| 630047 | 1 | East Asia | Makuwa | Korea | S | Ia-1 | VI-b | Pop B2 | 1.23 |
| 940147 | 1 | East Asia | Makuwa | Korea | S | Ib-1 | VI-b | Pop B2 | 1.45 |
| Takada-shiro-uri | 1 | East Asia | Conomon | Japan | S | Ia-1 | VI-b | Pop B2 | 2.12 |
| Tokyo-wase-shiro-uri | 1 | East Asia | Conomon | Japan | S | Ia-1 | VI-b | Pop B2 | 1.71 |
| Nakasaki-tsuke-uri | 1 | East Asia | Conomon | Japan | S | Ia-1 | VI-b | Pop B2 | 1.99 |
| Karimori | 1 | East Asia | Conomon | Japan | S | Ia-1 | VI-b | Pop B2 | 1.42 |
| Hyougo-aoshima-uri | 1 | East Asia | Conomon | Japan | S | Ia-1 | VI-b | Pop B2 | 1.90 |
| Kanro | 1 | East Asia | Makuwa | Japan | S | Ia-1 | VI-b | Pop B2 | 1.29 |
| Kinpyo | 1 | East Asia | Makuwa | Japan | S | Ia-1 | VI-b | Pop B2 | 1.28 |
| Seikan | 1 | East Asia | Makuwa | Japan | S | Ia-1 | VI-b | Pop B2 | 0.91 |
| Nanbukin | 1 | East Asia | Makuwa | Japan | S | Ia-1 | VI-b | Pop B2 | 1.79 |
| Kohimeuri | 3 | East Asia | Makuwa | Japan | S | Ia-1 | VI-b | Pop B2 | 0.80 |
| Weedy melon | 3 | East Asia | Agrestis | Korea | S | Ia-3 | VI-b | Pop B2 | 0.82 |
| 940082 | 1 | East Asia | Agrestis | Korea | S | Ia-3 | VI-b | Pop B2 | – |
| 940009 | 1 | East Asia | Agrestis | Japan | S | Ia-1 | VI-b | Pop B2 | 0.53 |
| 940012 | 1 | East Asia | Agrestis | Japan | S | Ia-1 | VI-b | Pop B2 | 0.56 |
| 940047 | 1 | East Asia | Agrestis | Japan | S | Ia-1 | VI-b | Pop B2 | 0.54 |
| 940111 | 1 | Western Africa | Agrestis | Sudan | S | Ic-5/-6 | IV | Pop A3 | – |
| PI 185111 | 2 | Western Africa | Agrestis | Ghana | S | Ic-1 | IV | Pop A1/A3 | – |
| 940108 | 1 | Western Africa | Agrestis | Senegal | S | Ic-3/-4 | IV | Pop B1 | – |
| 940112 | 1 | Western Africa | Agrestis | Senegal | S | Ic-3/-4 | IV | Pop B1 | – |
| PI 436532 | 2 | Western Africa | Agrestis | Senegal | S | Ic-3/-4 | IV | Pop A1/A3 | – |
| PI 436534 | 2 | Western Africa | Agrestis | Senegal | S | Ic-3/-4 | IV | Pop A3 | – |
| 940065 | 1 | Western Africa | Agrestis | Cameroon | S | Ic-5/-6 | IV | Pop A3 | – |
| 940107 | 1 | Southern Africa | Agrestis | South Africa | S | Ia-1 | VI-c | Pop B1 | – |
| PI 525105 | 2 | Northern Africa | Unclassified landrace | Egypt | L | Ib-1 | Ia | Pop A1 | – |
| PI 525109 | 2 | Northern Africa | Unclassified landrace | Egypt | L | Ib-2 | Ia | Pop A1 | – |
| PI 525110 | 2 | Northern Africa | Unclassified landrace | Egypt | L | Ib-2 | Ia | Pop A1 | – |
| PI 525111 | 2 | Northern Africa | Unclassified landrace | Egypt | L | Ib-2 | Ia | Pop A1 | – |
| PI 525114 | 2 | Northern Africa | Unclassified landrace | Egypt | L | Ib-2 | Ia | Pop A1 | – |
| PI 527386 | 2 | Northern Africa | Unclassified landrace | Algeria | L | Ib-3 | Ia | Pop A1 | – |
| PI 207658 | 2 | Northern Africa | Unclassified landrace | Morocco | L | Ib-1 | IIc | Pop A2 | – |
| PI 207661 | 2 | Northern Africa | Unclassified landrace | Morocco | L | Ib-3 | IIa | Pop A2 | – |
| PI 490388 | 2 | Western Africa | Unclassified landrace | Mali | L | Ib-2 | VII | Pop A2 | – |
| 940281 | 1 | Western Africa | Unclassified landrace | Chad | S | Ic-5/-6 | IV | Pop A3 | – |
| PI 436533 | 2 | Western Africa | Unclassified landrace | Senegal | S | Ic-2 | Ic | Pop A1/A3 | – |
| PI 320993 | 2 | Western Africa | Unclassified landrace | Sierra Leone | L | Ia-1 | Ib | Pop A1 | – |
| Cam-84-3 | 1 | Western Africa | Unclassified landrace | Cameroon | S | Ic-5/-6 | IV | Pop A3 | – |
| PI 505599 | 2 | Southern Africa | Unclassified landrace | Zambia | S | Ia-1 | IV | Pop A3 | – |
| PI 505602 | 2 | Southern Africa | Unclassified landrace | Zambia | S | Ia-1 | VI-c | Pop B1 | – |
| PI 482398 | 2 | Southern Africa | Unclassified landrace | Zimbabwe | S | Ic-5/-6 | IV | Pop A3 | – |
| PI 482411 | 2 | Southern Africa | Unclassified landrace | Zimbabwe | S | Ia-1 | VII | Pop B1 | – |
| PI 482413 | 2 | Southern Africa | Unclassified landrace | Zimbabwe | S | Ic-5/-6 | IV | Pop A3 | – |
| PI 482424 | 2 | Southern Africa | Unclassified landrace | Zimbabwe | S | Ia-3 | VII | Pop B1 | – |
| PI 482429 | 2 | Southern Africa | Unclassified landrace | Zimbabwe | S | Ia-3 | VII | Pop B1 | – |
| PI 234607 | 2 | Southern Africa | Unclassified landrace | South Africa | L | Ib-3 | Ic | Pop A1 | – |

[a] Two cultivars listed as ‘Ogen’ and ‘Cantaloure de Bellegrade’ are considered as ‘Ha'Ogen’ and ‘Cantaloup de Bellegarde’, respectively.

[b] Tendral o Invernale a Buccia Verde.

[c] Seed source is indicated by numbers: 1 = NARO Institute of Vegetable and Floriculture Science (NARO/NIVFS), Japan. 2 = North Central Regional Plant Introduction Station, Iowa State University (USDA-ARS), USA. 3 = Okayama University, Japan.

[d] S: Small seed-type, L: Large seed-type.

Supplemental Table 2. Accession number of nucleotide sequence registered in the DNA Data Bank of Japan

| Sequence accession number | Accession number | Cytoplasm type | Sequence region |
|---|---|---|---|
| LC822783 | 940102 | Ia-5 | Protein-coding region of *psbK* gene and *psbK*–*psbI* intergenic region |
| LC822784 | PI 614519 | Ia-6 | Protein-coding region of *psbK* gene and *psbK*–*psbI* intergenic region |
| LC822785 | 940102 | Ia-5 | Protein-coding region of *rpl16* gene and *rpl16*–*rpl14* inter-genic region (PS-ID) |
| LC822786 | PI 614519 | Ia-6 | Protein-coding region of *rpl16* gene and *rpl16*–*rpl14* inter-genic region (PS-ID) |
| LC822787 | 940099 | Ia-4 | Protein-coding region of *ndhA* gene and *ndhA* intron1 |
| LC822788 | 940102 | Ia-5 | Protein-coding region of *ndhA* gene and *ndhA* intron1 |

Supplemental Table 3. Statistics of the 27 RAPD markers generated from 212 melon accessions

| Marker name | Primer sequence (5' to 3') | Number of different alleles | Number of effective alleles | PIC |
|---|---|---|---|---|
| A07-1353 | GATGGATTTGGG | 2.0 | 1.9 | 0.126 |
| A07-872 | GATGGATTTGGG | 2.0 | 1.5 | 0.171 |
| A20-1100 | TTGCCGGGACCA | 2.0 | 2.0 | 0.196 |
| A20-800 | TTGCCGGGACCA | 2.0 | 1.3 | 0.044 |
| A22-1520 | TCCAAGCTACCA | 2.0 | 1.2 | 0.085 |
| A23-1200 | AAGTGGTGGTAT | 2.0 | 1.5 | 0.235 |
| A26-1400 | GGTGAGGATTCA | 2.0 | 1.5 | 0.126 |
| A31-800 | GGTGGTGGTATC | 2.0 | 2.0 | 0.229 |
| A39-2027 | CCTGAGGTAACT | 2.0 | 1.6 | 0.187 |
| A41-1353 | TGGTAGGTAACT | 2.0 | 1.5 | 0.196 |
| A41-1020 | TGGTAGGTAACT | 2.0 | 1.3 | 0.175 |
| A41-930 | TGGTAGGTAACT | 2.0 | 2.0 | 0.257 |
| A57-800 | ATCATTGGCGAA | 2.0 | 2.0 | 0.221 |
| B15-600 | CCTTGGCATCGG | 2.0 | 1.9 | 0.209 |
| B32-900 | ATCATCGTACGT | 2.0 | 2.0 | 0.252 |
| B32-700 | ATCATCGTACGT | 2.0 | 1.8 | 0.223 |
| B68-1078 | CACACTCGTCAT | 2.0 | 2.0 | 0.225 |
| B71-1220 | GGACCTCCATCG | 2.0 | 1.5 | 0.244 |
| B84-700 | CTTATGGATCCG | 2.0 | 1.4 | 0.180 |
| B84-600 | CTTATGGATCCG | 2.0 | 1.9 | 0.282 |
| B84-550 | CTTATGGATCCG | 2.0 | 1.3 | 0.123 |
| B86-1500 | ATCGAGCGAACG | 2.0 | 1.2 | 0.076 |
| B86-1350 | ATCGAGCGAACG | 2.0 | 1.6 | 0.114 |
| B96-850 | CTGAAGACTATG | 2.0 | 1.5 | 0.240 |
| B96-750 | CTGAAGACTATG | 2.0 | 1.9 | 0.230 |
| B99-1400 | TTCTGCTCGAAA | 2.0 | 1.6 | 0.156 |
| C00-1350 | GAGTTGTATGCG | 2.0 | 1.7 | 0.163 |

Supplemental Table 4. Classification of 130 melon accessions into large and small seed-types based on 100 seed weight

| Seed type | No. of accessions | | | 100 seed weight (g) | | |
|---|---|---|---|---|---|---|
| | Total | 2.2 g < | 2.2 g ≥ | Average | Minimum | Maximum |
| Large seed | 46 | 3 | 43 | 3.47 ± 1.16 | 2.03 | 8.03 |
| Small seed | 84 | 79 | 5 | 1.36 ± 0.57 | 0.36 | 2.73 |

Supplemental Table 5. Genetic variation in 10 groups of melon classified by geographical origin and seed size type.

| Area / Seed size | Number of accessions | Number of alleles | Number of effective alleles | Gene diversity |
|---|---|---|---|---|
| Europe/US_large seed | 28 | 1.78 | 1.54 | 0.261 |
| West and Central Asia_large seed | 22 | 1.89 | 1.58 | 0.318 |
| West and Central Asia_small seed | 3 | 1.19 | 1.32 | 0.230 |
| South Asia_large seed | 25 | 1.93 | 1.67 | 0.367 |
| South Asia_small seed | 48 | 2.00 | 1.52 | 0.292 |
| Southeast Asia_large seed | 6 | 1.67 | 1.50 | 0.307 |
| Southeast Asia_small seed | 17 | 1.78 | 1.35 | 0.211 |
| East Asia_small seed | 33 | 1.48 | 1.27 | 0.147 |
| Africa_large seed | 11 | 1.59 | 1.52 | 0.223 |
| Africa_small seed | 18 | 1.78 | 1.42 | 0.262 |

Supplemental Table 6. Pairwise estimates of $F_{ST}$ and genetic distance among five model-based subpopulations

| Subpopulation | PopA1 | PopA2 | PopA3 | PopB1 | PopB2 |
|---|---|---|---|---|---|
| PopA1 | - | 0.2144 | 0.5754 | 0.3974 | 0.6119 |
| PopA2 | 0.1343 | - | 0.5340 | 0.2404 | 0.5164 |
| PopA3 | 0.3528 | 0.3733 | - | 0.5948 | 0.6628 |
| PopB1 | 0.3054 | 0.1517 | 0.3227 | - | 0.3462 |
| PopB2 | 0.6156 | 0.3842 | 0.4550 | 0.1353 | - |

[a] $F_{ST}$ is shown above the diagonal, and pairwise genetic distance below the diagonal.
